# Mechanical Regulation of Stem Cell Proliferation and Fate Decisions by their Differentiated Daughters

**DOI:** 10.1101/2020.08.31.275081

**Authors:** Wenxiu Ning, Andrew Muroyama, Hua Li, Terry Lechler

## Abstract

Basal stem cells fuel development, homeostasis, and regeneration of the epidermis. The proliferation and fate decisions of these cells are highly regulated by their microenvironment, including the basement membrane and underlying mesenchymal cells. Basal progenitors give rise to differentiated progeny that serve an essential role in generating the epidermal barrier. Here, we present data that differentiated progeny also regulate the proliferation, differentiation, and migration of basal progenitor cells. Using two distinct mouse lines, we found that increasing contractility of differentiated cells resulted in non-cell autonomous hyperproliferation of stem cells and prevented their commitment to a hair follicle lineage. These phenotypes were rescued by pharmacological inhibitors of contractility. Live-imaging revealed that increasing the contractility of differentiated cells resulted in stabilization of adherens junctions and impaired movement of basal progenitors during hair placode morphogenesis, as well as a defect in migration of melanoblasts. These data suggest that intra-tissue tension regulates stem cell proliferation, fate decisions and migration, similar to the known roles of extracellular matrix rigidity. Additionally, this work demonstrates that differentiated epidermal keratinocytes are a component of the stem cell niche that regulates development and homeostasis of the skin.

## Introduction

Stem/progenitor cells drive the development, homeostasis and wound-healing of many tissues, including the epidermis^1^. In the skin, these progenitors lie in the basal layer of the epidermis, where they make contact with an underlying basement membrane that separates them from the underlying dermis (Fig. 1a). During skin development and homeostasis, the basal stem cells exhibit multiple cell behaviors including proliferation and differentiation to generate suprabasal cells, thus maintaining a functional barrier. In addition, during embryonic development and wound-healing, basal cells must migrate to aid in the formation of hair follicles or to heal cuts^2–4^. Understanding how stem cell fates and behaviors are determined by cues from their local microenvironment, termed the niche, is a fundamental goal to control and exploit stem cell activity.

**Fig. 1:**
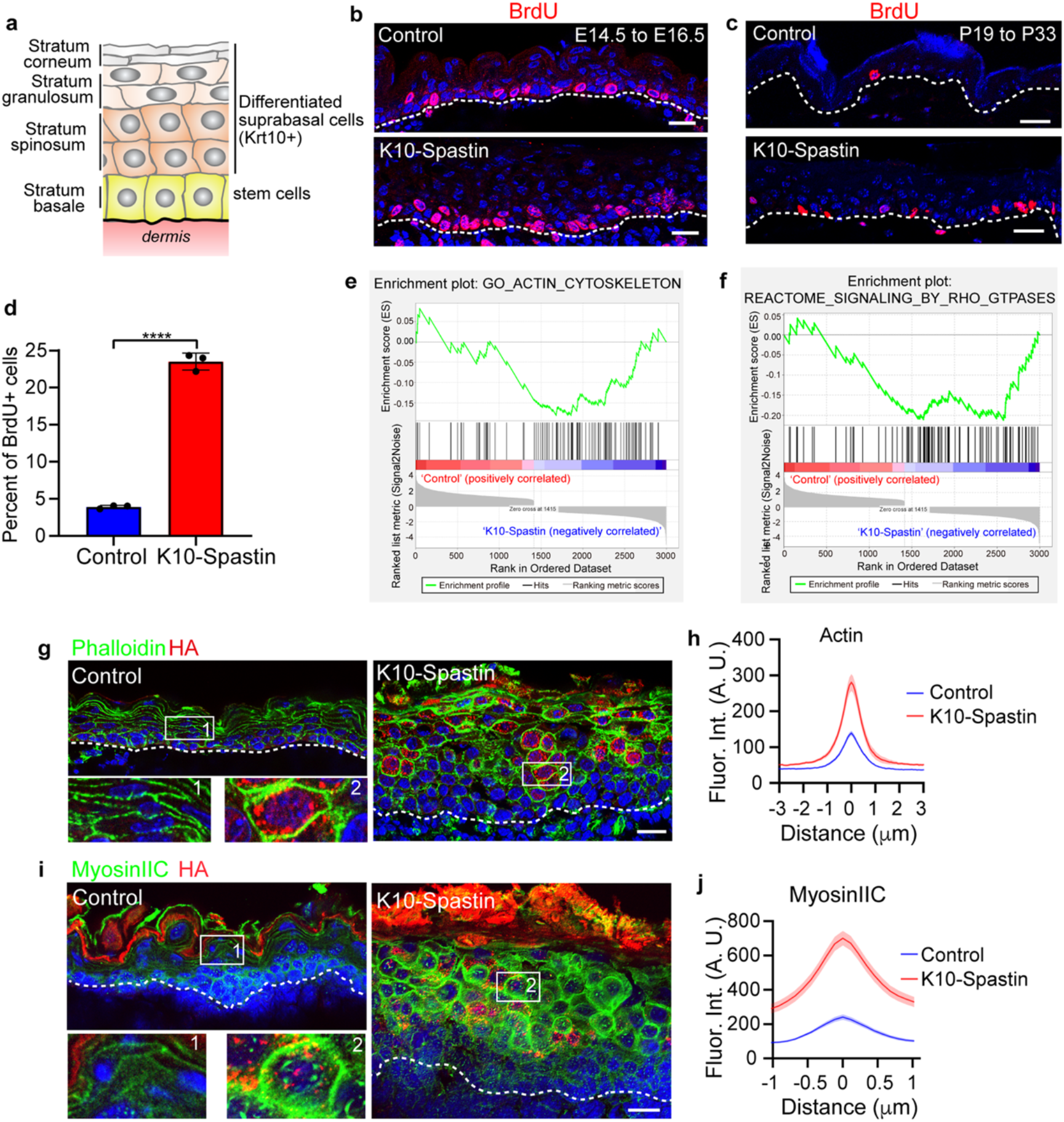
Microtubule disruption increases cortical actomyosin. **a**, Schematic depicting the stratified epidermis with basal stem cells and their progeny - Krt10+ differentiated suprabasal cells. **b-c**, BrdU (red) immunofluorescence staining in control and K10-Spastin epidermis at E16.5 (b), and in adult (c). Embryos/mice were exposed to doxycycline from E14.5-16.5 for (b) and from P19-33 for (c). Scale bars, 20 μm. **d**, Percentage of BrdU+ basal cells in adult treated skin epidermis. Data is shown as the mean ± SD, n=3 mice for control (36 fields) and K10-Spastin (38 fields) were measured, p-value<0.0001, two-tailed unpaired t-test. **e-f**, Gene Set Enrichment Analysis (GSEA) of RNA-Seq data for control and K10-Spastin, revealing of actin cytoskeleton gene signature (e) and Rho-GTPase reactome signaling related genes (f) enriched in K10-Spastin. Normalized enrichment score (NES): −1.42, FDR: 0.016, p-value: 0.003 for (e), and NES: −1.56, FDR: 0.0125, p-value: 0.003 for (f). **g**, Immunofluorescence staining of F-actin labeled by phalloidin (green) and Spastin-HA (red) in the epidermis at E16.5. Doxycycline treated from E14.5. Scale bar, 20 μm. Frame1 and 2 are enlarged control and spastin-expressing cells respectively. **h**, Quantification of F-actin fluorescence intensity at cell-cell boundaries of suprabasal cells in control and K10-Spastin. Data shown as the mean ± SEM, n=38 cells for control and n=39 cells for K10-Spastin from 3 embryos were measured, p-value <0.0001, two-tailed unpaired t-test. **i**, Immunofluorescence staining of cortical MyosinIIC (green) and Spastin-HA (red) at E16.5, doxycycline was added at E14.5. Scale bar, 20 μm. Frame1 and 2 are enlarged control and spastin-expressed cells respectively. **j**, Quantification of MyosinIIC fluorescence intensity at cell-cell boundaries of suprabasal cells in control and K10-Spastin. Data is shown as the mean ± SEM, n=40 cells for control and n=50 cells for K10-Spastin from 3 embryos, p-value <0.0001, two-tailed unpaired t-test.

Known niche factors in the epidermis include signals from the underlying dermis^5–7^. There are important roles for factors secreted from mesenchymal cells, including dermal fibroblasts, adipose cells, vasculature, neurons, immune cells and lymphatics, in regulating the behavior of stem cells in the skin^5, 8–12^. Notably, both basement membrane attachment and the rigidity of the underlying substrate play regulatory roles in stem cell proliferation/differentiation decisions^13, 14^. Basal stem cells are adhered to the extracellular matrix (ECM) via cell surface receptors, including integrins, which help organize ECM into the basement membrane. Integrin attachment and signaling are important to maintain proliferation and prevent differentiation of stem cells in the skin^15–19^. Stem cells are sensitive not only to integrin attachment, but also to the rigidity of the basement membrane and underlying dermis. Increased rigidity promotes the proliferation of basal stem cells, presumably through integrin sensing and mechanotransduction to intrinsic actin cytoskeleton dynamics, though the details of this have not been well deciphered in the skin^20–25^. Another possible niche signal is from the progeny of the stem cells. These differentiated daughters remain in physical contact with their stem cells, however, roles for these cells in regulating stem cells have not been reported.

Recently, paracrine signaling roles for differentiated daughters in controlling stem cell behavior have begun to be elucidated in other tissues^26, 27^. For example, in the *Drosophila* midgut, daughter enteroendocrine cells regulate Notch signaling in adjacent stem cells to prevent their differentiation, and enterocyte death causes tissue damage and compensatory increases in stem cell proliferation^28–31^. In the fly hematopoietic system, progeny release factors that maintain stem cell quiescence^32^. In the lung, injury induces expression of BMP antagonists, promoting proliferation of stem cells^33^. Unlike these tissues, the skin epidermis is stratified and has multiple cell layers (Fig. 1a). These cell layers are attached to each other by cell-cell adhesion structures, thus allowing for direct cell-cell communication. Therefore, they have the potential for both paracrine and direct mechanical signaling in both directions. However, the potential niche functions of differentiated cells, and their mechanical coupling to the basal progenitor have not been directly addressed.

Previously, we developed mouse lines to allow inducible gene expression in the differentiated cells of the epidermis^34^. The fast turnover of these cells has limited traditional recombination-based loss of function studies. Notably, we found that disruption of microtubules, specifically in differentiated cells of the epidermis, resulted in dramatic thickening of this tissue^34^. This was due to both cell-autonomous effects on cell shape in the differentiated cells and a non-cell autonomous increase in stem cell proliferation^34^. Here, we show that microtubule disruption results in increased acto-myosin contractility. Increased contractility is both necessary and sufficient for the phenotypes resulting from microtubule disruption. Increased contractility not only causes proliferation of basal progenitor cells, but also alters cell fates, - inhibiting cells from fully committing to hair follicles. These effects are mediated, at least in part, by a cell migration defect due to increased contractility of overlying cells. Melanoblasts, which also migrate among basal progenitor cells, have similar defects in their migration. Our data demonstrate an increase in cell-cell adhesion stability, which likely underlies these effects. Therefore, we propose that stem cells of stratified tissues not only respond to substrate rigidity through integrin signaling, but also to intra-tissue tension, at least partially through effects on adherens junctions. Further, this work establishes that the mechanical state of the differentiated cells is a niche signal that regulates multiple aspects of stem cell behavior.

## Results

### Microtubule disruption results in increased cortical actomyosin

We previously showed that we could efficiently disassemble microtubules in differentiated epidermis by driving expression of an active form of the microtubule severing protein, spastin (K10-rtTA;TRE-HA-spastin, hereafter referred to as K10-Spastin)^34^. Microtubule depolymerization in differentiated suprabasal cells of the epidermis leads to hyperproliferation of basal stem/progenitor cells in a non-cell autonomous manner (Fig. 1b-d, Fig. 3f). This effect occurred both during development and homeostasis (Fig. 1c-d), and did not depend on the barrier activity of the epidermis (i.e. it is not a simple response to loss of physiological function of the tissue) ^34^. To explore the underlying mechanism of how differentiated cells regulate skin development and stem cell proliferation, we isolated keratin 10-postive (K10^+ve^) differentiated cells from both control and K10-Spastin embryos. Cells were sorted based on the expression of H2B-GFP, driven by K10-rtTA. Embryos were treated with doxycycline at e14.5 to induce both spastin and H2B-GFP expression, then collected and FACS sorted for the GFP+ suprabasal cells at e16.5 (Extended Data Fig. 1a-b). Gene ontology (GO) term analysis of the resulting RNA-Seq dataset showed changes in gene signatures for epidermal development, peptide crosslinking, and lipid catabolism (Extended Data Fig. 1e). These are late epidermal differentiation signatures, consistent with our previous finding of premature barrier formation in spastin-expressing mice^34^. GO term and gene set enrichment analysis (GSEA) also revealed changes in actin cytoskeleton genes and in regulators of Rho-GTPase signaling (Fig 1e-f, Extended Data Fig. 1c, d). As the actin cytoskeleton is a major determinant of cell shape, this offered a potential explanation for the cell-autonomous changes in cell morphology in these mice. Further, we noted that many of the significant changes were in genes associated with actomyosin contractility. A ‘contractome’, containing just over 100 proteins that regulate acto-myosin contractility has been defined^35^, and we found that about 50% of these genes were differentially expressed in differentiated cells upon microtubule disruption, including myosin II-C, and the upstream factors Rock2, RhoC and Arhgef2 (Extended Data Fig. 1c). Thus, our RNAseq analysis suggested that microtubule disruption specifically in differentiated cells might alter F-actin organization and contractility.

To validate these findings, we first stained actin filaments in K10-Spastin embryos. Quantification of the intensity across intercellular junctions of the microtubule-disrupted cells showed a dramatic increase in cortical F-actin (Fig. 1g-h). Notably, F-actin level between basal cells and their contacted spastin-expressing spinous cells was also higher than in controls, demonstrating changes at this important interface (Extended Data Fig. 1f). Non-muscle myosin II molecules assemble into bipolar filaments and bind to F-actin to generate contractile forces^36^. Myosin IIC, which showed an approximately 12 fold increase at the mRNA level, also showed an enrichment at the protein level by western blot (Extended Data Fig. 1g). It was highly enriched at the cortex of the differentiated cells in K10-Spastin embryos (Fig. 1i-j). The increase in cortical Myosin IIC was cell autonomous and thus restricted to differentiated cells that expressed spastin (Fig. 1i). In contrast, myosin II A and IIB were not significantly altered in the mutant epidermis (Extended Data Fig. 1h-i). These data indicate that microtubule depolymerization can increase cortical actomyosin contractility in suprabasal differentiated cells of the skin epidermis.

### Microtubule disruption stabilizes adherens junctions

Cortical actomyosin complexes can interact with and productively engage adherens junctions. Adherens junctions are mechanosensitive and responsive structures that can be stabilized by the external application of force^37, 38^. Since cortical actomyosin was increased in microtubule disrupted suprabasal cells, we asked whether the increased cortical actomyosin altered adherens junctions *in vivo*. We therefore stained for with the α-18 antibody, which specifically recognizes an epitope of α-catenin that is exposed when junctions are under tension^38^. Staining and quantification revealed that there was not a significant change in the levels of total α-catenin at the cell cortex (Fig. 2a-b). However, cortical α-18 was increased in K10-Spastin as compared to control skin (Fig. 2a, c). Co-staining of α-18 and F-actin demonstrated that they were increased in the same cells (Fig. 2a).

**Fig. 2:**
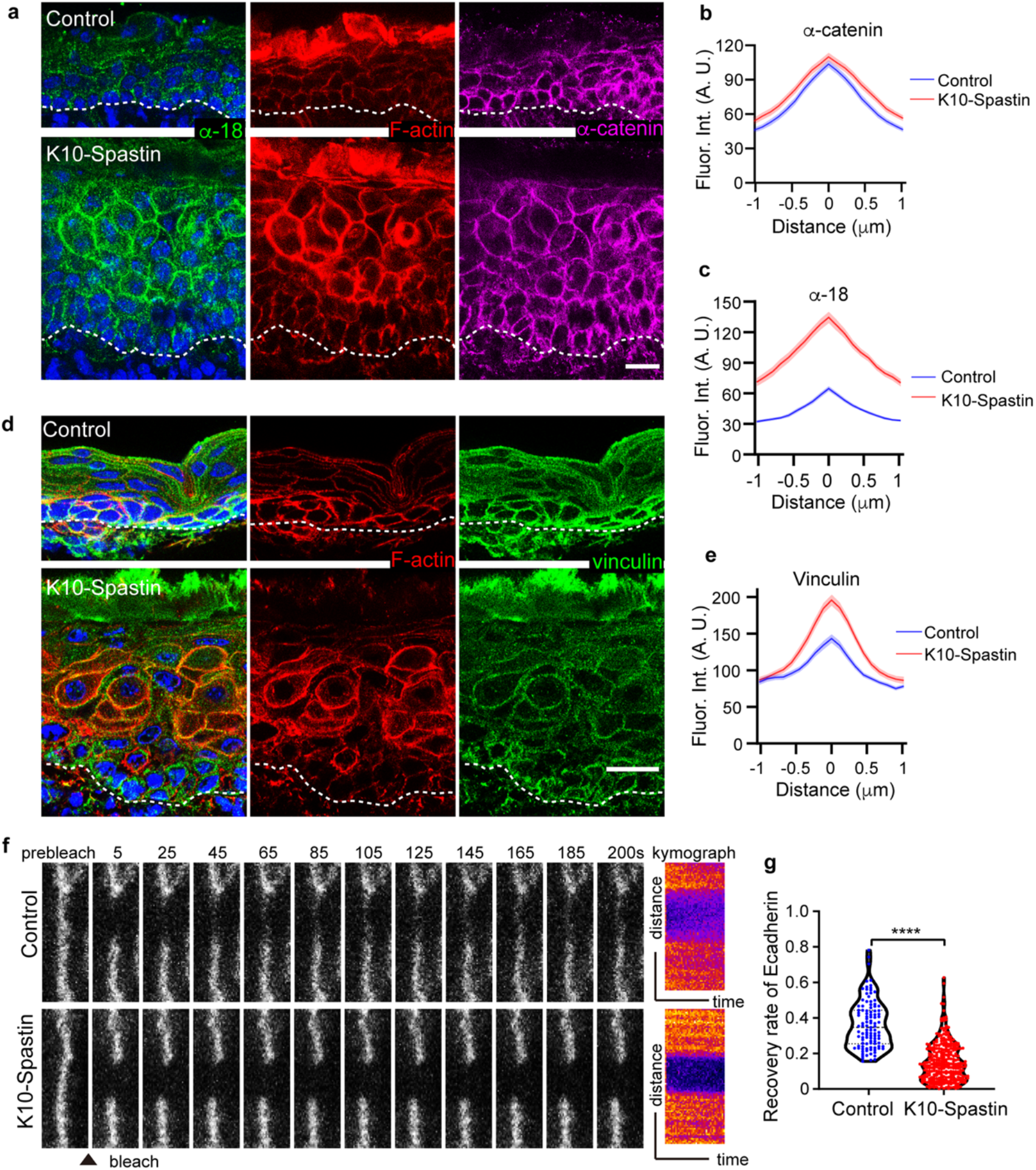
Microtubule depolymerization increases adherens junctional tension and stability. **a**, Immunofluorescence staining of α-18 (green), F-actin (red) and α-catenin (magenta) in control and K10-Spastin epidermis. Scale bar, 20 μm. **b**, Quantification of α-catenin fluorescence intensity at cell-cell boundaries of suprabasal cells in control and K10-Spastin epidermis. Data shown as the mean ± SEM, n=40 cells for control and n=41 cells for K10-Spastin from 3 embryos, p-value = 0.205, two-tailed unpaired t-test. **c**, Quantification of α-18 fluorescence intensity at cell-cell boundaries of suprabasal cells in control and K10-Spastin. Data shown as the mean ± SEM, n=35 cells for control and n=42 cells for K10-Spastin from 3 embryos, p-value <0.0001, two-tailed unpaired t-test. **d**, Co-staining of vinculin (green) and F-actin (red) in control and K10-Spastin epidermis. Scale bar, 20 μm. **e**, Quantification of vinculin fluorescence intensity at cell-cell boundaries of suprabasal cells in control and K10-Spastin epidermis. Data shown as the mean ± SEM, n=29 cells for control and K10-Spastin from 2 embryos, p-value <0.0001, two-tailed unpaired t-test. **f**, Time lapse series of E-cadherin-CFP FRAP in suprabasal cells in control and K10-Spastin epidermis at E15.5. Kymograph of E-cadherin-CFP over 200 seconds is shown on the right. **g**, Quantification of E-cadherin-CFP recovery rate in 200 seconds after photobleaching in suprabasal cells is shown in violin plot. For control, n=107 regions from 6 embryos, for K10-Spastin, n= 305 regions from 3 embryos, p-value<0.0001, two-tailed unpaired t-test.

In addition to the exposure of the α-18 epitope, vinculin often marks adherens junctions under increased tension^38^. The conformation of α-catenin that can productively bind to vinculin is the same that is recognized by α-18. Consistent with the increase in cortical α-18 signal, we observed increased cortical intensity of vinculin in microtubule disrupted cells (Fig. 2d-e). Another readout of contractility in many cell types is the mechano-responsive transcription factors YAP/TAZ. This protein is nuclear in many basal progenitors where it plays an important role in driving proliferation^39^. In differentiated cells of control epidermis, YAP is undetectable. However, there was a clear nuclear YAP signal in the differentiated epidermis of K10-Spastin embryos (Extended Data Fig. 2a-b). As most basal progenitors in the embryo are YAP positive in control mice, there was not a clear increase in the mutant. However, in the adult skin, YAP positive basal cells are rare in control epidermis, while they were increased in the K10-Spastin mutant (Extended Data Fig. 2c-e). Together, these data demonstrate that microtubule depolymerization increases actomyosin contractility thus engaging adherens junctions.

One effect of increased junctional tension can be a change in stability and/or turnover of E-cadherin complexes^40, 41^. To directly test whether spastin expression alters adherens junction dynamics, we used an E-cadherin-CFP mouse line crossed to control or spastin-expressing background. This is a fully functional E-cadherin that is tagged at its endogenous chromosomal locus^42^. Fluorescence recovery after photobleaching (FRAP) experiments were performed on skin explants. In normal epidermis, E-cadherin-CFP recovered to about 36% of its initial intensity within 200 seconds. In K10-Spastin epidermis, recovery was only 15% (Fig. 2f, g). These data demonstrate that the increased actomyosin contractility induced by microtubule disruption not only leads to increased junctional tension but also to stabilization of cortical E-cadherin complexes.

### Contractility is required for both cell autonomous and non-cell autonomous effects of microtubule disruption

While contractility was increased in spastin-expressing cells, it was unclear whether it was required to drive the cell autonomous effects (changes in cell shape of differentiated cells) or the non-cell autonomous effects (changes in stem cell proliferation) caused by microtubule depolymerization. We thus treated embryos with inhibitors of contractility to determine whether this could rescue either of the phenotypes. Non-muscle myosins were targeted with blebbistatin^43^, while the upstream regulator, ROCK1/2, was inhibited with Y-27632^44^. We injected pregnant dams with these inhibitors to decrease myosin activity (Extended Data Fig. 3d). Decreasing contractility with either of these inhibitors rescued the skin thickness as well as cell aspect ratio compared to untreated K10-Spastin embryos (Fig. 3a-e). Cell aspect ratio was not rescued completely, however, and it remains unclear whether this is because of incomplete inhibition or whether microtubules play a more direct role in cell shape control (Fig. 3d). Remarkably, decreasing contractility also rescued basal cell proliferation in K10-Spastin epidermis. The inhibitors did not have a significant effect on proliferation in control epidermis (Fig. 3f, Extended Data Fig. 3a-c).

**Fig. 3:**
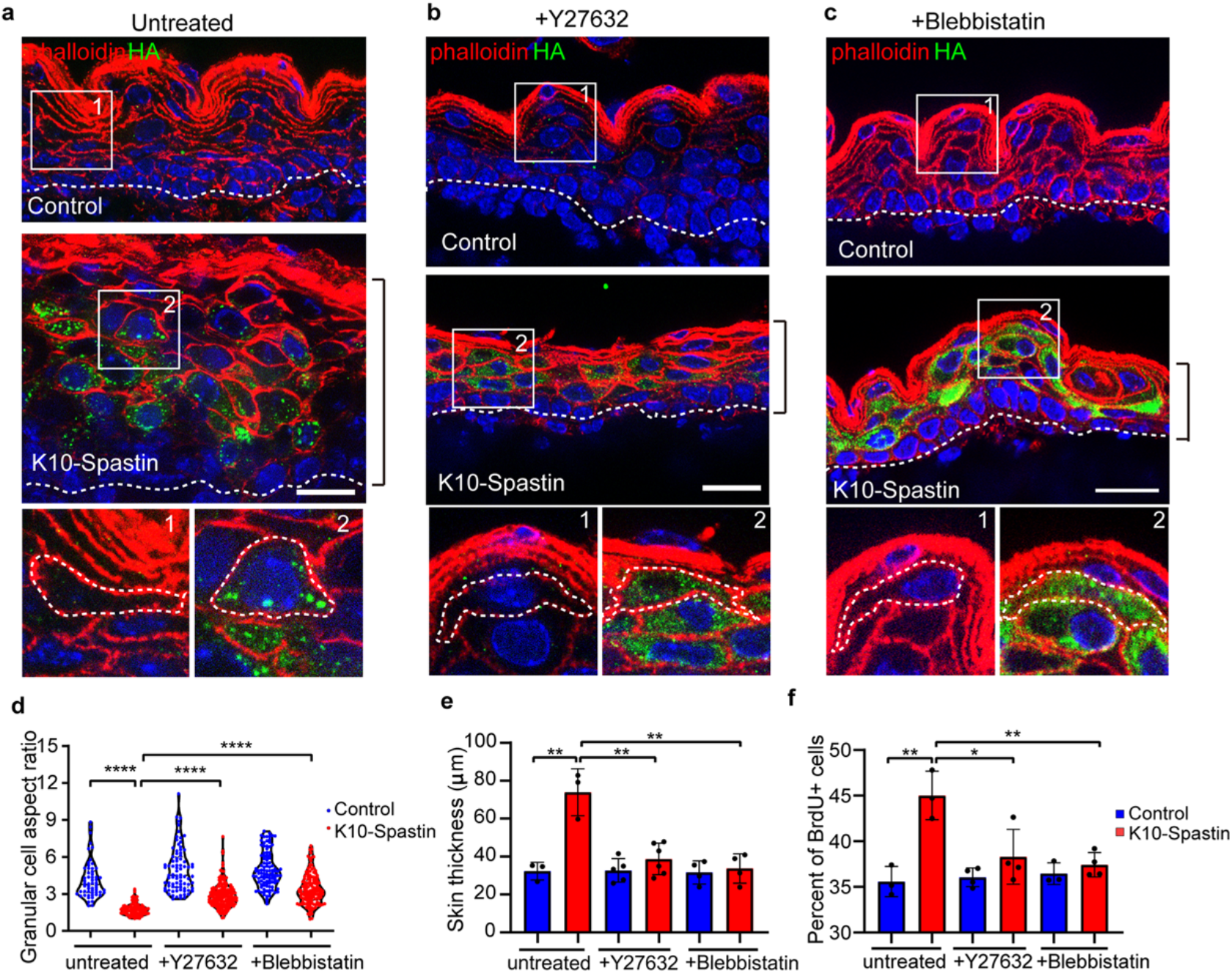
Decreasing contractility rescues skin thickness, cell shape and basal cell hyperproliferation caused by microtubule disruption in suprabasal cells. **a**, **b**, **c**, Immunofluorescence staining of Spastin-HA (green) and F-actin (red) in untreated, Y27632, and Blebbistatin injected control and K10-Spastin epidermis respectively. For each condition, frames 1 and 2 are enlarged cells from control and K10-Spastin respectively. Scale bars: 20 μm. **d**, Quantification of granular cell aspect ratio in untreated, Y27632, and Blebbistatin treated control and K10-Spastin epidermis. Data is shown as the mean ± SD. For the untreated condition, n=58 cells for control and n=102 cells for K10-Spatin from 3 embryos. For Y27632, n=83 cells for control and n=171 cells for K10-Spastin from 4 embryos. For blebbistatin, n=105 cells for control and n=146 cells for K10-Spastin from 4 embryos. For the analysis of the labeled group: p-value<0.0001, two-tailed unpaired t-test. **e**, Quantification of skin thickness in untreated, Y27632, and Blebbistatin treated control and K10-Spastin epidermis. Data is shown as the mean ± SD. For untreated condition, n=3 embryos for control (89 regions) and K10-Spastin (102 regions) from 2 untreated litters were measured. For Y27632, n=5 embryos for control (182 regions) and n=6 embryos for K10-Spastin (255 regions) from 3 injected litters were measured. For blebbistatin, n=4 embryos for control (133 regions) and K10-Spastin (214 regions) from 3 injected litters were measured. For the analysis of the labeled group: p-value<0.01, two-tailed unpaired t-test. **f**, Percentage of BrdU+ basal cells in untreated, Y27632, and Blebbistatin treated control and K10-Spastin epidermis. Data is shown as the mean ± SD. For untreated, n=3 embryos for control (31 fields) and K10-Spastin (40 fields). For Y27632, n=4 embryos for control (32 fields) and K10-Spastin (38 fields). For blebbistatin, n=3 embryos for control (31 fields) and n=4 embryos for K10-Spastin (47 fields). For the analysis: control to K10-Spastin untreated group: p-value<0.01; K10-Spastin untreated to Y27632 treated group: p-value=0.0279; K10-Spastin untreated to blebbistatin treated group: p-value<0.01; all are two-tailed unpaired t-test.

### Contractility of differentiated daughters is sufficient to induce stem cell proliferation

The above data demonstrate that contractility of differentiated cells is necessary to induce progenitor cell proliferation, but does not address whether it is sufficient. To further test a direct role for suprabasal cell contractility in regulating basal stem cells and skin development, we developed a mouse line that inducibly increases contractility using a constitutively active Rho-GEF (TRE-HA-Arhgef11^CA^) to increase actomyosin contractility without disrupting microtubules. Arhgef11 is a PDZ-domain-containing Rho guanine nucleotide exchange factor (RhoGEF) that can bind and activate RhoA through its catalytic DH-PH domains^45^ (Fig. 4a). Transfection of this construct with K14-rtTA into cultured keratinocytes revealed membrane localization of Arhgef11^CA^ and cell rounding with increased cortical F-actin (Extended Data Fig. 4a). Notably, neighbor cells were elongated toward the contracting Arhgef11^CA^ cells, consistent with increased contractility.

**Fig. 4:**
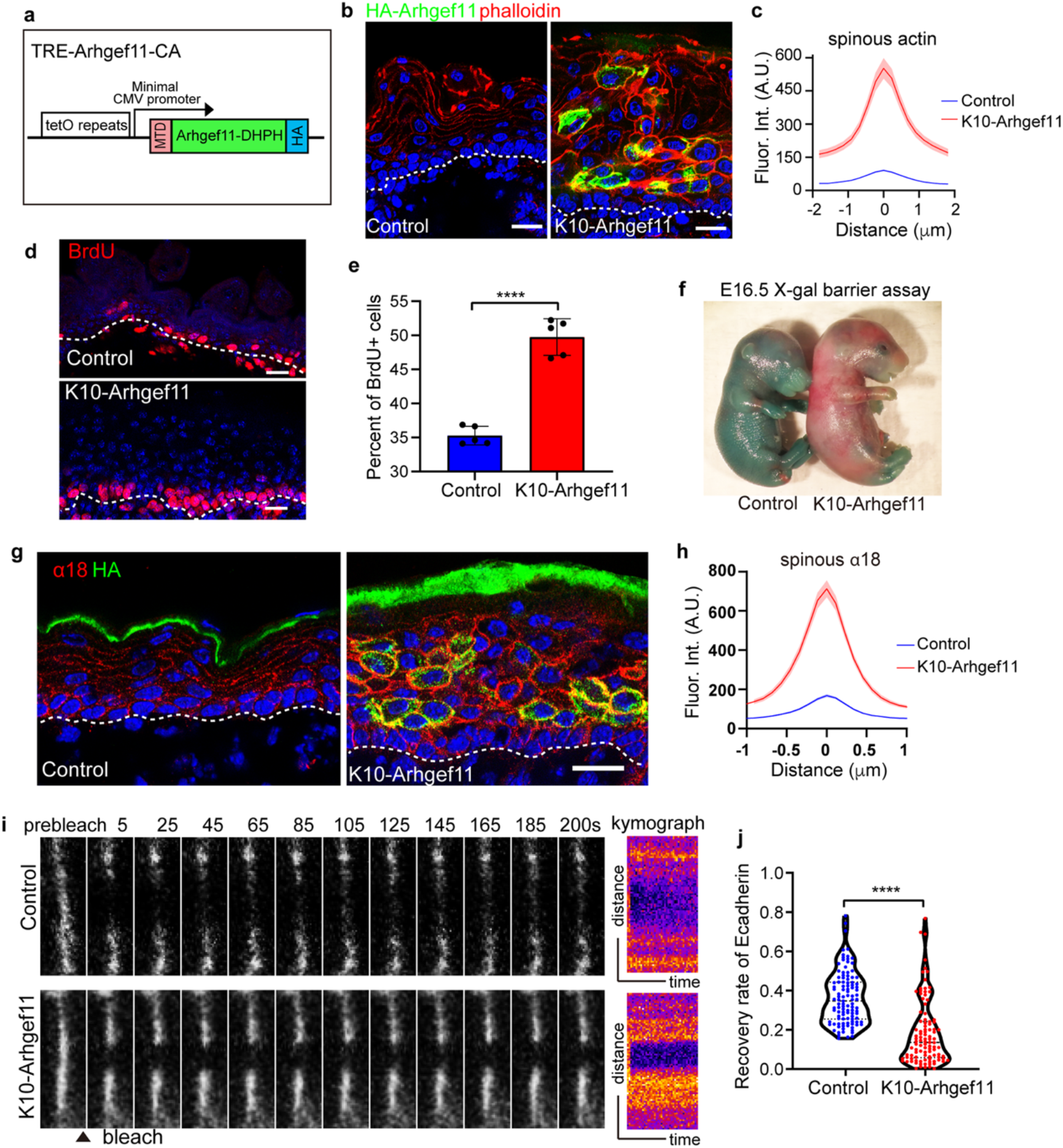
RHO-GEF induced contractility stabilizes adherens junction and induces stem cell proliferation. **a**, Diagram of the TRE-Arhgef11^CA^ transgene. Arhgef11-DHPH was tagged with a membrane targeting domain (MTD) at the N-terminus and an HA tag at the C-terminus. **b**, Co-staining of F-actin (phalloidin in red) and Arhgef11-HA (green) in the epidermis of control and K10-Arhgef11 mice. Scale bar, 20 μm. **c**, Quantification of cortical F-actin fluorescence intensity in control and K10-Arhgef11spinous cells. Data shown as the mean ± SEM, n=27 cells for control and n=33 cells for K10-Arhgef11 from 3 embryos, p-value <0.0001, two-tailed unpaired t-test. **d**, BrdU staining in control and K10-Arhgef11 epidermis at E17.5, doxycycline-treated from E13.5. Scale bar, 20 μm. **e**, Percentage of BrdU+ basal cells in control and K10-Arhgef11 epidermis. Data is shown as the mean ± SD. n=5 embryos for control (48 fields) and K10-Arhgef11 (40 fields), p-value<0.0001, two-tailed unpaired t-test. **f**, Skin X-gal barrier assay of e16.5 embryos from control and K10-Arhgef11 embryos. Absence of dye indicates an effective outside-in barrier. **g**, Co-staining of α-18 (red) and Arhgef11-HA (green) in control and K10-Arhgef11 epidermis at E17.5. The HA staining has non-specific auto-fluorescence signal in the cornified layer. Scale bar, 20 μm. **h**, Quantification of cortical α-18 fluorescence intensity in control and K10-Arhgef11spinous cells. Data shown as the mean ± SEM, n=35 cells for control and n=51 cells for K10-Arhgef11 from 3 embryos, p-value <0.0001, two-tailed unpaired t-test. **i**, Time lapse series of E-cadherin-CFP FRAP in suprabasal cells in control and K10-Arhgef11epidermis at E15.5. Kymograph of E-cadherin-CFP over 200 seconds is shown on the right. **j**, Quantification of E-cadherin-CFP recovery rate in 200 seconds after photobleaching in suprabasal cells. Note that the dataset displayed for the control here is the same as in Fig. 2f-g. For control, n=107 regions from 6 embryos, for K10-Arhgef11, n= 102 regions from 3 embryos, p-value<0.0001, two-tailed unpaired t-test.

We established a mouse line K10rtTA; TRE-Arhgef11^CA^ (hereafter referred to as K10-Arhgef11) that allows doxycycline-inducible Arhgef11^CA^ expression in suprabasal differentiated cells, and found a similar cortical localization in the differentiated cells *in vivo* as we observed in cultured keratinocytes *in vitro* (Fig. 4a-b). Grossly, the mice expressing Arhgef11^CA^ in suprabasal cells presented with thick flaky skin, lack of external ears, misshapen paws, and curly tails (Extended Data Fig. 4c). In addition, these mice had the same premature barrier acquisition phenotype and impaired corneocyte morphology as observed in K10-Spastin mice (Fig. 4f, Extended Data Fig. 4k). There was a clear increase in tissue contractility when Arhgef11^CA^ was induced in *ex vivo* cultured skin, resulting in a shrinking of the entire skin over time (Extended Data Fig. 4b).

These similarities were also reflected at the cell and tissue levels. K10-Arhgef11 embryos had a thickened epidermis, decreased aspect ratio of granular and spinous cells, and an increase in progenitor cell proliferation (Fig. 4d-e, Extended Data Fig. 4h-l). In addition, Arhgef11^CA^ expression increased cortical F-actin and α-18 staining intensity both in the granular cells and spinous cells, similar to the effects of spastin (Fig. 4b-c, g-h, Extended Data Fig. 4d-e). Notably, the intensity of α-18 staining at the interface between basal progenitors and Arhgef11^CA^ expressing spinous cells was also increased compared to controls, reflecting the increased junctional tension across this interface (Extended Data Fig. 4f-g). To confirm whether junctional stability was increased by the Arhgef11^CA^-expressing tissue, we performed FRAP analysis of E-cadherin-CFP in these embryos. This revealed a similar level of adhesion strengthening as was observed in spastin-expressing embryos (Fig. 4i-j, Fig. 2f-g). Together, these data demonstrate that increased contractility of differentiated cells is both necessary and sufficient to induce stem cell proliferation.

To determine the spatial localization of contractility that is required to promote stem cell proliferation, we turned to another driver line, Involucrin-tTA, that allows doxycycline to repress expression from the TRE promoter (i.e. a tet-off system, in contrast to the tet-on provided by rtTA) in upper spinous and granular cells of the epidermis, but is absent from immediate suprabasal cells^46^ (Extended Data Fig. 5a). In this system, doxycycline was removed to induce protein expression. In combination with the TRE-Arhgef11 allele, we found mosaic expression throughout the epidermis at different developmental stages. As expected, at E16.5, Arhgef11^CA^ expression was largely restricted to the upper spinous and granular layers. However, in some focal areas it was found even in the spinous cells immediately adjacent to the basal progenitors (Extended Data Fig. 5a-b). At E18.5, Arhgef11^CA^ was more robustly expressed in the first layer of spinous cells as well as the suprabasal cells of hair follicles (Extended Data Fig. 5e). We therefore quantitated basal proliferation at E16.5 in areas where there were fewer contacted spinous cells expressing Arhgef11^CA^, and areas where there were more. We found that with increasing numbers of adjacent cells expressing Arhgef11^CA^, there were increasing levels of basal progenitor cell proliferation (Extended Data Fig. 5b-d). These data are consistent with direct cell-cell contact being required between the contractile cell and the basal progenitor, and are inconsistent with this signal being transmitted across multiple cells or systemically.

### Contractility causes non-cell autonomous defects in hair follicle formation

During skin development, the epidermal stem cells in the basal layer undergo fate decisions. They either differentiate and move up into stratified interfollicular epidermis or invaginate and migrate towards the dermis to generate hair follicles. When we induced spastin expression at e16.5, the resulting pups had less hair (Fig. 5a-b). Hair follicle development occurs in three consecutive waves, with the first wave starting around E14.5 forming guard hairs, the second wave starting at E16.5 forming awl/auchene hairs, and the third wave starting at E18.5 forming zigzag hairs^47^. To understand whether hair follicle development could be affected by the increased actomyosin contractility of Krt10+ suprabasal cells, we induced expression of spastin or Arhgef11^CA^ at E14.5 and analyzed hair follicles at later stages. Lhx2, a transcription factor that is expressed in hair follicles beginning at the late placode stage, was used to quantitate hair follicle number. There was a clear decline in hair follicles in both mouse models (Fig. 5c-f). To begin to determine the temporal window in which contractility works, we also induced expression at later time points, E16.5 and E18.5, and examined effects at postnatal day zero (P0 - neonates) (Extended Data Fig 6a-d). The later the treatment of doxycycline, the less severe the impact on hair follicle numbers in neonatal mice, suggesting that contractility likely acted early during hair follicle morphogenesis and did not significantly affect hair follicles once they had passed some threshold stage of development. As the first wave of hair placodes is already specified at E14.5, concomitant with the induction of expression of the spastin or Arhgef11^CA^, these hair follicles are likely spared from their effects. That said, we cannot rule out that the guard hairs are less sensitive to these effects.

**Fig. 5:**
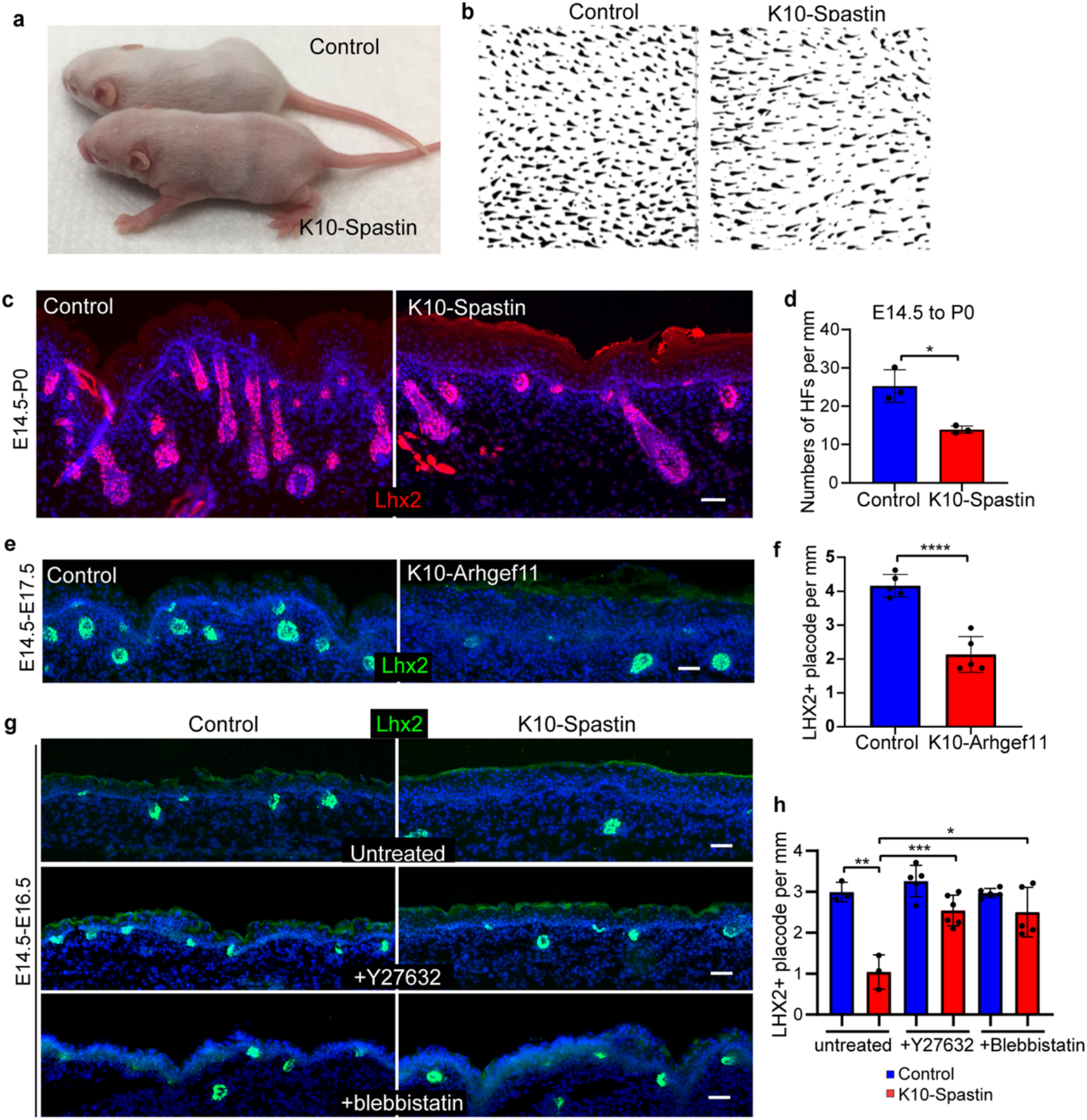
Increased contractility in differentiated cells causes non-cell autonomous defects in hair follicles formation. **a**, Decrease in hairs in K10-Spastin mice compared to controls at postnatal day 8. Spastin expression was induced at e16.5 with doxycycline. **b,**Whole mount of hair follicles in K10-Spastin and control mouse. **c-d,**Staining of LHX2 (red) to mark hair follicles in control and epidermis at P0. Expression was induced at E14.5. Scale bar, 50 μm. Data shown as the mean ± SD, n=3 embryos for control (24 fields) and K10-Spastin (31 fields), p-value= 0.0108, two-tailed unpaired t-test. **e-f,**Staining of LHX2 (green) in control and K10-Arhgef11 epidermis at E17.5. Expression was induced at E14.5. Scale bar, 50 μm. Data shown as the mean ± SD, n=5 embryos for control (62 fields) and K10-Arhgef11 (52 fields), p-value <0.0001, two-tailed unpaired t-test. **g-h,**Staining of LHX2 (green) in untreated, Y27632 or blebbistatin treated control and K10-Spastin embryos at E16.5. Expression was induced at E14.5. Scale bar, 50 μm. Data is shown as the mean ± SD. For untreated, n=3 embryos for control (36 fields) and K10-Spastin (46 fields). For Y27632, n=5 embryos for control (60 fields) and n=6 embryos for K10-Spastin (68 fields). For blebbistatin, n=5 embryos for control (65 fields) and K10-Spastin (70 fields). For the analysis: control to K10-Spastin untreated: p-value<0.01; K10-Spastin untreated to Y27632 treated: p-value<0.001; K10-Spastin untreated to blebbistatin treated: p-value= 0.0107; two-tailed unpaired t-test.

The hair follicle defects seen in K10-Spastin embryos were largely rescued by decreasing contractility with Y27632 and blebbistatin (Fig. 5g, h). Consistently, directly increasing actomyosin contractility by expressing Arhgef11^CA^ also decreased hair follicle specification (Fig. 5e, f).

Hair follicle morphogenesis begins with the formation of a thickened epithelium, called a placode (Fig. 6a). Wnt signaling is required for hair follicle specification^48^, and Lef1 marks cells as they commit to the hair fate (note that it also marks underlying dermal condensate cells). Subsequently, additional regulators, first EDAR, and then Lhx2 are expressed. In the late hair placode, Lhx2 labels the basal cells, while Sox9+ marks a population of cells directly above them^49^. To verify that the phenotypes we observed were in fact non-cell autonomous, we examined the position of Krt10^+ve^ cells relative to hair follicle placodes using K10-rtTA;TRE-H2B-GFP embryos. At the earliest stages of placode formation, H2B-GFP was expressed in cells overlying the forming placode, but not in the cells that are committing to a placode fate (Extended Data fig. 6i). At slightly later stages, a suprabasal layer of placode cells forms and contains a population of cells that are Sox9^+ve^. These cells did not express H2B-GFP and we never detected spastin or Arhgef11^CA^ in these cells (Extended Data fig. 6j). This indicates that the hair follicle specification defects in these two mouse models resulted from non-cell autonomous effects on actomyosin contractility in overlying differentiated cells. This is striking as no prior reports have shown that suprabasal cells can regulate basal stem cell fate during hair follicle specification.

**Fig. 6:**
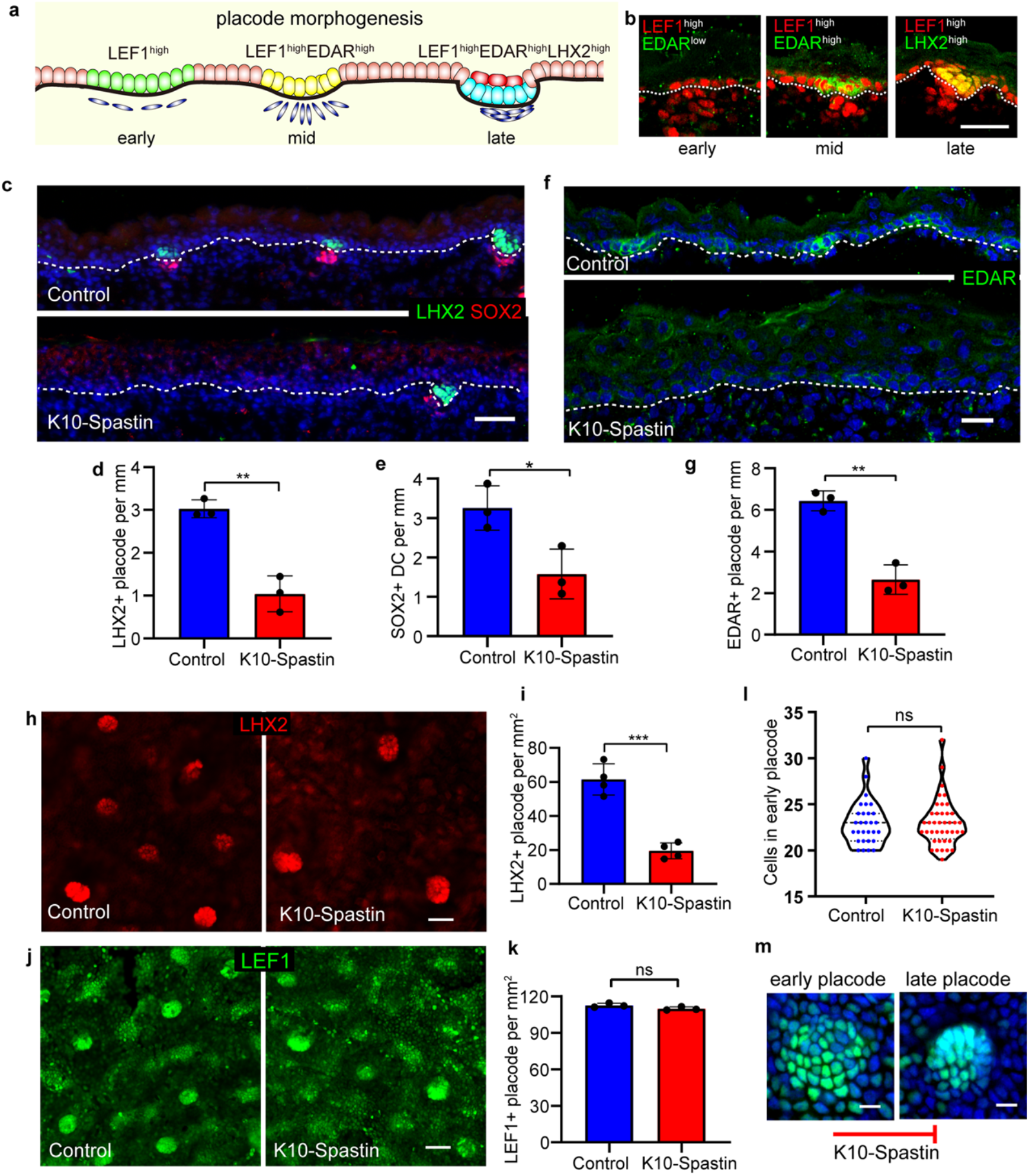
Increasing contractility in differentiated cells inhibits placode progression but not specification. **a**, Diagram depicting stages of hair placode by LEF1/EDAR/LHX2. **b**, Immunostaining of Lef1^high^ (red) early placode, EDAR^high^ mid placode, Lhx2^high^ late placode at E16.5. Scale bar, 30 μm. **c**, Co-staining of Lhx2(green) and Sox2(red) in control and K10-Spastin skin at E16.5. Expression induced at E14.5. Scale bar, 50 μm. **d**, Quantitation of Lhx2+ placodes in control and K10-Spastin E16.5 embryos. The data source is the same as in Fig. 5h. Data is shown as the mean ± SD. n=3 embryos for control (36 fields) and K10-Spastin (46 fields), p-value<0.01, two-tailed unpaired t-test. **e**, Quantification of Sox2+ dermal condensates (DC) in control and K10-Spastin embryos. Data is shown as the mean ± SD, n=3 embryos for control (36 fields) and K10-Spastin (41 fields), p-value=0.0266, two-tailed unpaired t-test. **f-g**, Decreased numbers of EDAR+ placodes in K10-Spastin compared to control at E16.5. Expression was induced at E14.5. Scale bar, 20 μm. Data shown as the mean ± SD, n=3 embryos for control (51 fields) and for K10-Spastin (61 fields), p-value<0.01, two-tailed unpaired t-test. **h-i,**Decreased Lhx2+ placodes in K10-Spastin compared to control by whole mount staining of the epidermis at E16.5. Expression induced at E14.5. Scale bar, 40 μm. Data shown as the mean ± SD, n=4 embryos for control (27 fields) and for K10-Spastin (28 fields), p-value<0.001, two-tailed unpaired t-test. **j-k,**Quantification of LEF+ placodes (including early and late placode) in control and K10-Spastin embryos at E16.5. Expression was induced at E14.5. Scale bar, 40 μm. Data shown as the mean ± SD, n=3 embryos for control (13 fields) and K10-Spastin (19 fields), p-value=0.1054 (>0.05, not significant), two-tailed unpaired t-test. **l,**Quantification of Lef1+ cell numbers in early placodes of control and K10-Spastin embryos at E16.5. Data shown as the mean ± SD, n=29 placodes for control and n=40 placodes for K10-Spastin from 3 embryos, p-value= 0.9546 (>0.05, not significant), two-tailed unpaired t-test. **m,**Images of Lef1 labeled early and late placode, the progression of which is blocked in K10-Spastin embryos. Scale bar, 10 μm.

### Contractility affects placode progression but not specification

Hair placode specification is initiated by Wnt/β-catenin signaling which is required for subsequent EDAR (Ectodysplasin receptor) expression^50, 51^. EDA/EDAR/NF-κB signaling is required to maintain and refine the pattern of Wnt/β-catenin activity at later stages of placode development^50^. Meanwhile, NF-κB directly targets *LHX2* which is required for the subsequent down-growth of the hair placode^52^. Therefore, in this study, we define Lef1, Edar, and Lhx2 expression as marking early, mid, and late placode stages (Fig. 6a-b, Extended Data fig. 6e-g). Consistently, the number of Lhx2^+ve^ placodes, as well as the number of dermal condensates (marked by Sox2), was decreased when microtubules were disrupted in differentiated cells (Fig. 6c-e).

To further explore earlier stages of hair placode formation, we stained for EDAR, which is required for LHX2 expression. The number of EDAR^+ve^ placodes was also lower in K10-Spastin embryos compared to controls (Fig. 6f-g, Extended Data Fig. 6h). We also stained for Lef1, the earliest marker of commitment to a hair follicle fate. This analysis was performed on whole-mount epidermis, which gives a clearer view of placode coalescence. The number of Lhx2^+ve^ placodes was decreased in K10-Spastin epidermis (Fig. 6h-i), consistent with our findings in sections. In control epidermis, early placodes were marked by Lef1 and were not highly condensed. Later stage placodes (also marked by Lhx2) were morphologically distinct, being tightly packed with clear boundaries (Fig. 6m). The total number of Lef1 patches was similar in the control and K10-Spastin mutant epidermis, demonstrating that initial hair follicle specification is not affected. However, the placodes in the K10-Spastin epidermis were mostly of the early/diffuse sort, with few of the more advanced placodes (Fig. 6j-k). The Lef1^+ve^ placodes contained similar numbers of cells in the control and mutant skin, demonstrating that subsequent defects were not due to a change in placode cell number (Fig. 6l). Therefore, suprabasal contractility regulates hair follicle fate downstream of their specification, and is required for both morphological and gene expression changes (Fig. 6m). Together, these data demonstrate that differentiated suprabasal cells not only regulate basal stem cell proliferation but also their cell fate during hair placode morphogenesis.

### Contractility decreases cell migration of placodal cells and melanoblasts

Prior work suggested cell migration drives placode coalescence during hair follicle initiation^2, 53^. We thus turned to live imaging of placode formation in skin explants to determine whether there were changes in cell dynamics in the K10-Spastin embryos. Using a membrane-bound GFP reporter, we observed placode cell dynamics in control and K10-Spastin embryos. In control embryos we found that early placodes underwent a consistent change in morphology, acquiring a smaller diameter over time. This change was not observed in the mutant skin (Fig. 7a-b). When quantitated subjectively by blinded observers, approximately 80% of placodes progressed in control skin, while less than 20% did in the mutant skin (Fig. 7c). We next tracked cell movements within the placode by labeling them with a K14-Cre; Rosa-LSL-H2B-mCherry reporter. Individual cell tracks were consistently shorter in the mutant as compared to the wild-type tissue, consistent with a defect in these cells’ ability to coalesce into a tight placode (Fig. 7d). Collectively, the live imaging results suggest that actomyosin contractility in differentiated suprabasal cells controls progenitor cell fates by regulating their cell migration during hair placode morphogenesis.

**Fig. 7:**
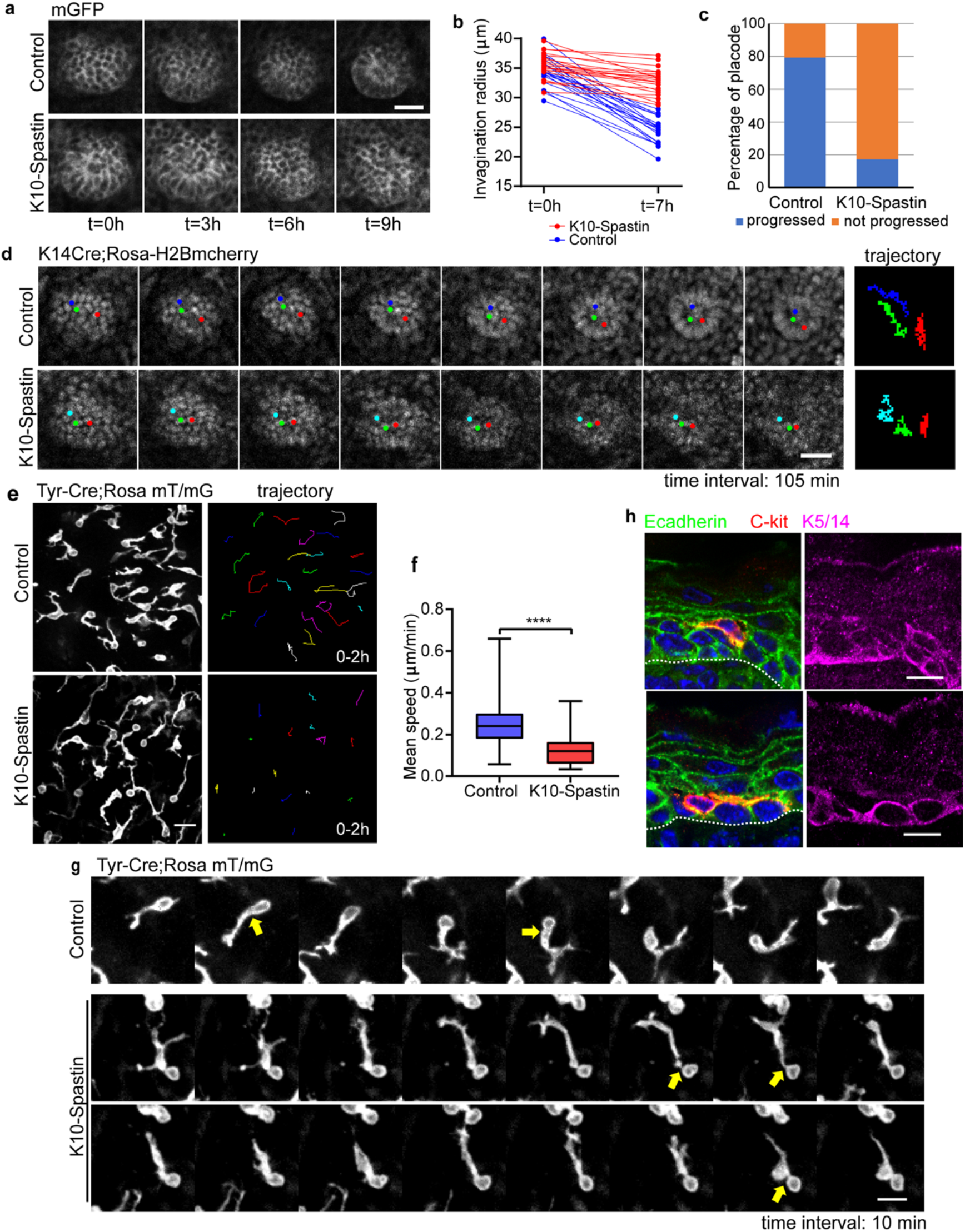
Increased contractility of differentiated epidermal cells decreases cell migration of placodal cells and melanoblasts. **a**, Time-lapse series of placode progression in control and K10-Spastin epidermis at E15.5, cell membranes are marked with a membrane-GFP. Scale bar, 20 μm. **b**, Quantification of placode radius in control and K10-Spastin skin at the beginning and the end of 7 hours imaging. n=20 placodes for control and n=22 placodes for K10-Spastin from 3 embryos. For ratio of radius[(0h-7h)/0h] between control and K10-Spastin, p-value<0.0001, two-tailed unpaired t-test. **c**, Percentage of placodes that progress in control and K10-Spastin embryos, n=58 placodes for control and n=46 placodes for K10-Spastin from 3 embryos. **d**, Time-lapse series of H2B-mCherry labeled placode progress in control and K10-Spastin epidermis imaged at E15.5. Time interval: 105 minutes, scale bar: 20 μm. **e**, Image of melanoblasts, as labeled by Tyr-Cre;Rosa mT/mG, in control and K10-Spastin skin at E15.5. The image on the right traces the movement of cells over a 2 hour imaging period. Scale bar, 20 μm. **f**, Quantification of the mean speed of melanoblasts in control and K10-Spastin epidermis. Data shown as the mean ± SD, n=484 melanoblasts from 4 embryos of control, and n=572 melanoblasts from 3 embryos of K10-Spastin were measured using Trackmate plugin in FIJI, p-value<0.0001, two-tailed unpaired t-test. **g**, Time lapse series of individual melanoblast migration in control and K10-Spastin epidermis imaged at E15.5. Arrows indicates the broader and larger protrusion next to cell body in control melanoblasts, while it is narrow and small in the melanoblast in the K10-Spastin skin. Scale bar: 10 μm, time interval: 10 minutes. **h**, Co-staining of E-cadherin (green), the melanoblast marker C-kit (red) and the basal cell marker (Krt5/14) in control epidermis at 16.5. Notice melanoblasts are stained with E-cadherin and exhibit migratory tendency through the interface between basal and spinous cell layers. Scale bar, 10 μm.

The embryonic epidermis consists not only of epidermal stem cells and their descendants, but also melanoblasts. Melanoblasts are neural-crest derived cells that invade into the epidermis at e12.5 and migrate quickly through the basal layer of progenitor cells, eventually homing to hair follicles in the mouse^54^. In tissue sections, staining of the melanoblast marker c-kit in the control epidermis demonstrated that the majority of melanoblasts sit within the basal layer of the epidermis, forming protrusions that traverse through lateral junctions, just above the basement membrane, and at the interface between suprabasal cells and basal cells (Fig. 7h, Extended Data Fig. 7d). To determine whether the effects of increased suprabasal contractility were unique to the keratinocyte lineage, we imaged the movement of melanoblasts that were labeled with Tyr-Cre; Rosa mT/mG (Fig. 7e). Melanoblasts migrated at an average speed of ~0.23 μm per minute in the control epidermis, and this was decreased to ~0.11 μm per minute in K10-Spastin embryos (Fig. 7e, f). Melanoblasts exhibited very dynamic protrusions from the cell body. Large protrusions were conduits through which the cell body could move (Fig. 7g). While K10-Spastin mice formed protrusions normally, the translocation of the cell body through these protrusions was rare (Fig. 7g). In support of this, the aspect ratio of the melanoblast cell body decreased significantly in K10-spastin, likely as a result of decreased motion and therefore decreased deformation as it squeezed between cells (Extended Data Fig. 7a). While melanoblasts in K10-Spastin embryos had normal numbers of protrusions, there was a slight increase in the length of the leading process, likely due to the lack of cell body movement into them (Extended Data Fig. 7b-c). This defect is consistent with the adhesion strengthening as junctional remodeling during migration is required.

Consistent with high expression of E-cadherin in migrating melanoblasts^54^, co-staining of E-cadherin with the melanoblast marker c-kit showed that melanoblasts form contacts with both basal and suprabasal keratinocytes, and have protrusions at the interface between basal and suprabasal layers (Fig. 7h, Extended Data Fig. 7d). Therefore, the melanoblast migration defects may result from direct effects on stabilized junctions between melanoblasts and keratinocytes or between cells through which they are traversing. F-actin and α-18 were increased at the interface between contracting spinous and basal cells (Extended Data Fig. 1f, 4g-h), though lateral F-actin and α-18 of basal keratinocytes were unchanged (Fig. 1e, 2a, 4b, 4g). Unfortunately, E-cadherin-CFP was too dim to perform FRAP analysis in basal cells to determine whether junctions are also stabilized at sites not in direct contact with the contractile cell.

In summary, these data demonstrate that differentiated epidermal cells play broad and unexpected roles in the skin. In addition to their role in generating a physiological barrier, they regulate the proliferation and fate decisions of underlying basal stem cells, as well as the migration of melanoblasts.

## Discussion

In this work, we demonstrated that the differentiated cells of the epidermis can regulate proliferation, migration and cell fate decisions of the underlying stem cells from which they are derived. Further, our evidence demonstrates that the contractility of these cells is key in determining stem cell behavior by both promoting proliferation and inhibiting migration.

A key finding in this work is that loss of microtubules results in increased cell contractility. Microtubule-actin crosstalk is mediated through both direct coupling by cross-linkers and by indirect signaling mechanisms that control Rho GTPases and their regulating GEFs and GAPs^55, 56^. In some other cell types, Arhgef2/GEF-H1, which is an activator for the contractility-inducing RhoA GTPase, can mediate this effect. This GEF has low activity when bound to microtubules, and that activity increases when microtubules are disrupted^57, 58^. However, we also saw a transcriptional upregulation of contractome genes. Therefore, it will be important to determine what mediates the transcriptional response and whether it is directly responsible for the contractility or a feedback mechanism that amplifies it. Notably, we find that this cellular response to microtubule disruption is not universal, as microtubule disruption in differentiated cells of the intestinal epithelium did not result in a similar contractility phenotype^59^. Consistent with this, the upregulation of contractility-related genes was not noted upon microtubule disruption in cultured retinal pigment epithelium cells, suggesting distinct responses of different cell types to microtubule disruption^60^. Further understanding of the mechanism will also inform how sensitive this response is to changes in microtubule dynamics and/or organization.

Regardless of the mechanism, it is clear that increased contractility of differentiated cells has profound effects on stem cell behavior. Importantly, these effects are not simply a response to a loss of tissue functions (i.e. sensing a barrier defect), as these mice have an intact barrier, and the effects are seen even before a barrier forms during embryogenesis. There are a number of ways in which contractility could regulate stem cell behavior. First, the effects could be indirect. Contractility in differentiated cells may alter their secretome or cell surface protein composition, thus resulting in local and or systemic changes (Extended Fig. 8). As the proliferation changes described here occurred locally, we can rule out systemic signaling. It is more difficult to determine the effects of changes to paracrine factors or direct contact of signaling molecules. However, a second intriguing possibility is that contractility acts via mechanical signaling to alter the behavior of underlying stem cells (Extended Data Fig. 8). In support of this possibility, we show that increased contractility results in changes in the dynamics of E-cadherin, a major transmembrane component of the adherens junctions in these cells. The significant stabilization of adherens junctions may contribute to the observed effects. In terms of migration, effective migration of cells through the basal layer requires the making and breaking of cell contacts. Decreased turnover of adherens junction would limit the ability to productively change neighbors and thus move within the tissue. Second, the direct application of force on adherens junctions in cultured cells was sufficient to induce their proliferation through both Yap1 and β-catenin-dependent pathways^61^. A similar effect in the skin may underlie the ability of differentiated cells to communicate mechanically with the underlying progenitors. In support of this, we saw an increase in nuclear YAP-positive basal cells when contractility was increased in differentiated cells in the adult epidermis. In contrast, we did not see an upregulation of Wnt target genes. It is notable that effects on actin dynamics and adherens junction turnover on one side of the junction are usually associated with changes on the other side, so it is likely that basal cells sense the changes in their overlying progeny^62^. In contrast, decreasing contractility in the epidermis does not appear to have significant effects on stem cells, as loss of RhoA or type II myosins did not have strong phenotypes^63–65^.

Epidermal stem cells sense and react to the mechanical stress from the niche, including the stiffness of the basement membrane/dermis. Growing epidermal cells *in vitro* on substrates with different stiffnesses affects their proliferation and differentiation, with stiff substrates stimulating epidermal cell proliferation and migration while inhibiting differentiation^22^. Unlike cell-ECM interactions, the role of intra-tissue contractility in regulating stem cell behavior has not been studied in detail. The data presented here is consistent with a similar mode of mechanosensation at adherens junctions that modified stem cell behavior.

Increasing contractility in differentiated cells had non-cell autonomous effects not only on basal cell proliferation but also on hair follicle formation. This defect stemmed from a defective migration of cells fated toward hair follicles. While the ultimate fate of these cells is unclear, we have not noted apoptotic bodies. One likely possibility is that, in the absence of clustering, the signals required to maintain their fates are lost and these cells revert into interfollicular stem cells. Similarly, the dermal condensates underlying the placodes were lost. Migration defects were not limited to basal stem cells, but also extended to melanoblasts. Melanoblasts migrate quickly through the relatively static basal stem cells. Transient adherens junctions form between melanoblasts and keratinocytes, and adhesions must be remodeled to allow the melanoblast nucleus to move through. It remains unclear if the relevant junctions affected here are between the differentiated cell and the melanoblast, or whether basal progenitor adhesions to each other are also stabilized, thus inhibiting melanoblast movement. Unfortunately, the E-cadherin-CFP signal is too dim in basal cells to directly test this possibility. In either case, this finding provides a novel perspective to explore contributions of the interaction between melanocytes and keratinocytes during metastatic melanoma formation. It also raises the question of whether the migration of other immune regulatory cells in the epidermis is affected by altered contractility of differentiated cells.

Finally, the work here has important implications for the analysis of gene function in stem cells. Traditional methods of genetic analysis of gene function in stem cells rely on tissue or cell-type specific recombination approaches. This technology obligately results in loss of gene function in the progeny of the stem cell as well. This makes it difficult to determine cell autonomous effects of gene function in stem cells from non-cell autonomous functions in their differentiated progeny. For example, prior studies on contractility regulators may misinterpret direct effects on stem cells, versus secondary effects due to changes in their progeny^20^. Therefore, new tools that are able to specifically cause changes in either basal progenitors or their progeny are required.

## Material and Methods

### Mouse models

All animal work was approved by Duke University’s Institutional Animal Care and Use Committee. All mice were maintained in a barrier facility with 12 hour light/dark cycles. Mouse strains used in this research were: K10-rtTA and TRE-Spastin mice were generated in the lab as previously described^34^. Involucrin-rTA mouse were previously described (Julie Segre publication). All other mice were purchased from Jackson Laboratories, and their stock number is indicated. TRE-H2B-GFP (005104); Rosa-mT/mG (007576); mGFP were generated by mating Rosa-mT/mG mice with a CMV-Cre-deleter strain; Tyr-Cre (029788); K14Cre (005107); Rosa26-LSL-H2B mCherry (023139); E-cadherin-CFP (016933). CD1 mice were purchased from Charles River (Strain code: 022).

### Plasmids

Arhgef11^CA^ with a membrane targeting domain and an HA tag with a stop code on the C-terminal was synthesized by GenScript. NheI and Sal I sites were inserted on the 5’ and 3’ ends of the Arhgef11^CA^ cassette, respectively and used to clone into the pTRE2 vector. Doxycycline-dependent expression of the Arhgef11 cassette was first verified in cultured keratinocytes by co-transfecting with K14-rtTA plasmid. The Arhgef11^CA^ plasmid was linearized with AatII and BsaxI, purified and was used by the Duke Transgenic Core to generate transgenic TRE-Arhgef11^CA^ mice via pronuclear injection.

### Inhibitor Experiments

For BrdU experiments, BrdU (10 mg/kg) was intraperitoneally injected into adult or pregnant dams (for embryos) one hour before sacrificed for tissue dissection and processing.

For drug rescue experiments in K10-Spastin, pregnant dams were fed with doxycycline chow starting at E14.5. They were intraperitoneally injected twice the next day (one early in the morning and one at night) with Y-27632 2HCl (Selleckchem, Cat. No. S1049) at 10 mg/kg per mouse, or with blebbistatin (Sigma, B0560) at 2 mg/kg per mouse.

### Cell culture

Stable wildtype keratinocytes established from primary mouse keratinoyctes were maintained in E low Ca^2+^ medium at 37°C and 7.5% CO_2_. Plasmids of K14-rtTA and pTRE2-Arhgef11^CA^ were co-transfected into keratinocytes using Mirus transfection reagents (Mirus). 12 hours after infection, cells were switched to E high Ca^2+^ medium (1.2 mM Ca^2+)^ to induce differentiation and with doxycycline (2 μg/ml) to induce Arhgef11^CA^ expression.

### Skin explant culture

Embryos at E15.5 were sacrificed, then rinsed and placed in sterile PBS. Back skin was cut off, rinsed, and gently unfolded in sterile PBS, then placed on an agarose pad (4% agarose in sterile water mixed 1:1 with 10%FBS in DMEM (Gibco, Cat. No.11965), and with doxycyline (2 μg/ml)), and 1:100 Penicillin/Streptomycin (Gibco, Cat. No.10378016)). Skin explants on agarose were gently placed into a 6 cm dish containing 1ml 10% FBS in Gibco-DMEM (to avoid agarose drying), and were cultured at 37°C and 7.5% CO_2_ for at least 2 hours for recovery before live imaging.

### RNA-Seq analysis

K10-rtTA;TRE-H2B-GFP;TRE-Spastin pregnant dams were fed with doxycycline chow at E14.5, then sacrificed at E16.5. Embryos were checked by dissecting microscope for GFP expression, tails were taken to confirm genotype, and back skins were cut off and treated with dispase II (1.4 U/ml) in HBSS at room temperature for 1 hour. The epidermis was then peeled off from the dermis, and digested in 1:1 trypsin (Gibco, Cat. No. 25200-056) with versene (Gibco, Cat. No. 15040-066) at 37°C and rotated for 20 minutes, then mixed 1:1 with FACS buffer (HBSS with 2.5% FBS and 10 μg/ml DNAaseI) and centrifuged. The cell pellet was diluted into FACS buffer with propidium iodide solution (Sigma P4864), filtered using sterile Celltrics 30 μm filters (04-004-2326, Sysmex), and FACS sorted for GFP positive and PI negative cells. RNA was extracted using a Qiagen RNAeasy Mini kit (Qiagen, Cat No.74104) following the manufacturer’s protocols, with DNA digested using RNase-Free DNase (Qiagen, Cat No.79254). Three independent RNA samples from control and K10-Spastin were collected and sent for sequencing and analysis by Novogene.

Differentially-expressed genes (p-value<0.01, FDR<0.05) by FPKM for certain pathway were centered, log2-transformed, and used to generate heatmaps. The complete differentially expressed genes (p-value<0.01, FDR<0.05, 2999 genes in total) were analyzed for the gene set enrichment using GSEA software^66^ (download and follow the guidelines from https://www.gsea-msigdb.org/gsea/index.jsp). GO term analysis for differentially expressed genes (fold-change>2 in K10-Spastin/control, p-value<0.01, FDR<0.05) was performed using WebGestalt^67^ online resources (http://www.webgestalt.org/).

### X-Gal barrier assay

E16.5 embryos were rinsed in PBS, then were placed into the staining solution (1 mg/ml X-gal, 1.3 mM MgCl_2_, 100 mM NaH_2_PO_4_, 3 mM K_3_Fe[CN]_6_, 3 mM K_4_Fe(CN)_6_, 0.01% sodium-deoxycholate, 0.2% NP-40), incubated at 37°C until color developed, washed in PBS for 1–2 minutes and then photographed.

### Cornified envelope preparations

Epidermis was isolated as above and then boiled for 10 minutes in 10 mM Tris (pH 7.4), 1% β-mercaptoethanol, and 1% SDS. Corneocytes were pelleted and resuspended in PBS and photographed.

### Western blot

Skin epidermal samples were prepared as previous described^65^. Epidermal proteins from control and K10-Spastin were solubilized in loading buffer (10% SDS, 40% Glycerol, 3% Bromophenyl Blue, and 10% β-mercaptoethanol), boiled for 10 minutes and loaded into 10% polyacrylamide gels and run for ~90 mins at 120V, then transferred onto nitrocellulose membrane, blocked with 5% BSA, and immunoblotted with primary antibodies for Myosin IIC (Biolegend, 919201) and GAPDH (Abcam, ab9485) overnight. Blots were washed three times in PBST (0.1% Tween in PBS), incubated with secondary antibodies (Licor, IRDye 680RD Series, CW800 Series), then visualized using a LI-COR Odyssey FC system.

### Immunofluorescence analysis

Skin tissue sections were fixed with 4% PFA at room temperature for 10 minutes or with ice-cold acetone (for Myosin IIC and vinculin) for 2 minutes, washed in PBS containing 0.2% Triton X-100, then incubated with blocking buffer (3% BSA with 5% NGS and 5% NDS for most antibodies, NGS was absent when using goat antibodies and MOM block was used for mouse primary antibodies). Sections were incubated in primary antibody diluted in blocking buffer for 1h at room temperature (α-18 incubated for 15 minutes), then washed, and incubated in secondary antibodies and Hoechst for 1 hour, washed, and finally mounted in the anti-fade buffer (90% glycerol in PBS plus 2.5 mg/ml p-Phenylenediamine (Sigma-Aldrich)).

For whole mount immunofluorescence staining, back skins were dissected off of embryos and treated with dispase at room temperature for 1 hour, and then the epidermis was peeled off from the dermis and fixed with 4% PFA for 30 minutes at room temperature. Skins were then washed and incubated with blocking buffer for 1 hour, and incubated with diluted primary antibodies in blocking buffer at 4 °C overnight. The next day they were washed again and incubated with secondary antibodies and Hoechst for 2 hours at room temperature, then washed and imaged.

Primary antibodies used in this study: rab anti-HA (Abcam, ab9110), rat anti-HA (Sigma-Aldrich, 11867423001), chicken anti-keratin 5/14 (generated in the Lechler lab), rabbit anti-keratin 10 (Covance, 905401), rat anti-BrdU (Abcam, ab6326), rat anti-β4 integrin (BD Biosciences, 553745), MyosinIIA (Biolegend, PRB-440P), MyosinIIB (Biolegend, 909901), MyosinIIC (Biolegend, 919201), α-catenin (Sigma-Aldrich, C2081), α-18 (gift from Akira Nagafuchi), Vinculin (Sigma-Aldrich, V9131), YAP/TAZ (Cell Signalling Technology, 8418S), LHX2 (Santa Cruz, 19344), SOX9 (Millipore, Ab5535), EDAR (Novus Bio, AF745), LEF1 (Cell Signaling Tech, 2230), E-cadherin (BD, 610182), c-Kit (Cell Signaling Technology, 3074S). F-actin was stained with phalloidin (Invitrogen, A12379 and Sigma-Aldrich, P1951).

### Imaging

For section and whole mount staining, slides were imaged on a Zeiss AxioImager Z1 microscope with Apotome.2 attachment, Plan-APOCHROMAT 20X/0.8 objective, Plan-NEOFLUAR 40X/1.3 oil objective, or Plan-NEOFLUAR 63X/1.4 oil objective, Axiocam 506 mono camera, and acquired using Zen software (Zeiss).

For live-imaging of hair placode and melanoblast migration, skin explants were placed upside down in a Lumox® dish 35 (94.6077.331, Sarstedt), with epidermal side facing toward the membrane. Images were acquired on an Andor XD revolution spinning disc confocal microscope at 37°C and 5% CO2 using a 20x/0.5 UplanFl N dry objective, and were acquired using MetaMorph software.

For FRAP assay of E-cadherin-CFP, skin explants were placed upside down on glass-bottom dishes (MatTek), with epidermis side facing toward the glass-bottom. FRAP assays were performed within one hour of mounting for each sample, using Leica SP5 Inverted Confocal microscope with 100x/1.4-0.70 oil objective, and was acquired using Leica confocal LAS AF software by setting up bleaching regions across junction, and taking time series before and after photobleaching.

### Image quantification and Statistics

All images quantifications were done using FIJI software. Quantifications of fluorescence intensity of cortical F-actin/Myosin IIC/α-catenin/α-18 were measured by drawing lines across cell-cell boundaries, and analyzed for their plot profiles, maxima were aligned and the ends were trimmed to yield the final line scan. Quantification of BrdU+ basal cells were performed by measuring numbers of BrdU+ cells and total basal cells for each field, and calculating the mean percentage of all fields for each mouse. Aspect ratios for granular and spinous cells were calculated by tracing individual cells and using the measurement option in FIJI to measure cell length and width. For recovery rate of E-cadherin-CFP, intensity was first normalized to the initial intensity before bleaching, then the fluorescence intensity was measured at time of prebleaching (I_0_), right after photobleaching (I_1_), and at 200 seconds after photobleaching (I_200_), and recovery rate= (I_200_ – I_1_)/(I_0_−I_1_). For melanoblasts, the number of protrusions generated from the cell body was counted manually, and the lengths of leading protrusion of melanoblasts were measured using FIJI. For quantification of melanoblasts migrating speed, the live imaging series were analyzed autonomously using the Trackmate^68^ plugin in FIJI to generate the mean speed.

All statistical analysis was performed using GraphPad Prism 5 software and Microsoft Excel. Statistical parameters including the exact value of n, meant ± SEM or ± SD, and statistical significance were mentioned individually in their relative figure legend. Data were judged to be statistically significant when p-value < 0.05 by two-tailed paired or unpaired Student’s t-test, asterisks denote statistical significance (ns = not significant, *, p < 0.05; **, p < 0.01, ***, p < 0.001, ****, p < 0.0001), as described in individual figure legend.

### Contributions

W.N. and T.L. conceptualized the study, designed the experiments, interpreted the data and wrote and reviewed the manuscript. W.N. performed most experiments, collected and quantified all the data. A.M. performed some hair follicle experiments in K10-Spastin. H.L. helped with GSEA and GO term analysis, and RNAseq heatmaps. T.L. acquired funding, supervised, and administered the project.

## Acknowledgements

We thank Julie Underwood for care of the mice and Elizabeth McDonald for help in genotyping. In addition, we thank Brent Hoffman and members of the Lechler Lab for comments on the manuscript. This work would not have been possible without reagents provided by Satrajit Sinha, Elaine Fuchs and Akira Nagafuchi. We thank the Duke Transgenic Core for the generation of TRE-arhgef11^CA^ mouse line, Bin Li from the Duke Flow Cytometry Shared Resource for cell sorting assistance, Yasheng Gao from the Duke Light Microscopy Core facility for imaging assistance. This work was supported by the Duke Regeneration Next Initiative (W.N), and by grants from the NIH to T.L. (R01-AR055926 and R01-AR067203).

---

**Extended Data Fig. 1:**
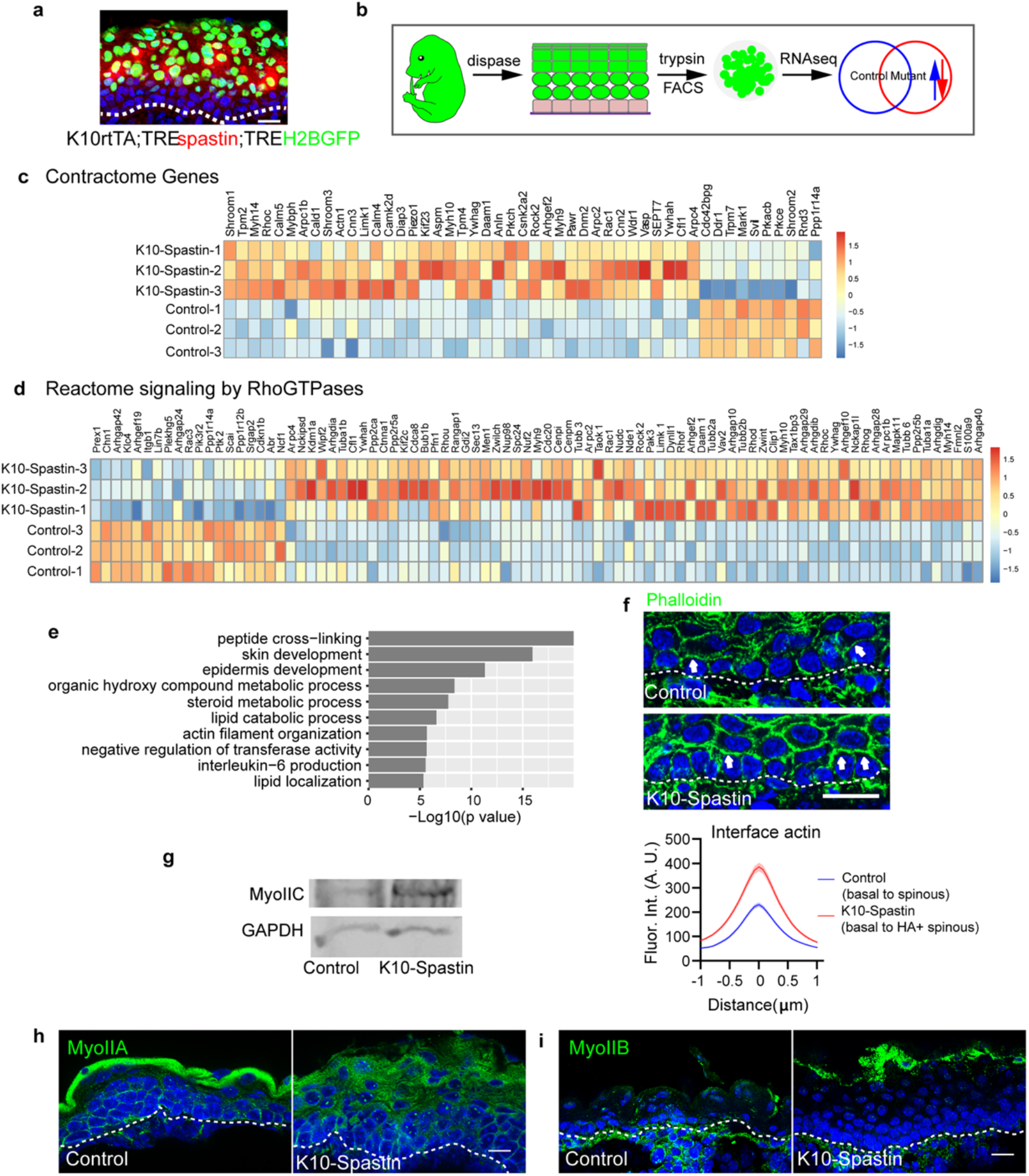
Microtubule disruption increases cortical actomyosin, related to Fig. 1. **a**, Immunostaining of HA labeled spastin (red) in K10rtTA;TREspastin;TRE-H2B-GFP, showing co-expression of spastin with H2B-GFP in suprabasal cells at E16.5. Scale bar, 20 μm. **b**, Diagram of RNA-Seq sample preparation. Only H2B-GFP labeled Krt10+ suprabasal cells from control and K10-Spastin were sorted and RNA isolated. n= 3 embryos for each group. **c**, RNA-Seq heatmap depicting differential expression of contractome genes in K10-Spastin and control suprabasal cells, gene expression by FPKM is centred and log2-transformed. **d**, RNA-Seq heatmap depicting differential expression of the Rho-GTPase reactome signaling genes in K10-Spastin and control suprabasal cells, gene expression by FPKM is centred and log2-transformed. **e**, Gene Ontology (GO) term enrichment analysis of genes (fold-change>2, p<0.01, FDR<0.05) for K10-Spastin compared to control, revealing upregulation of skin development and actin filament organization related biology process upon microtubule disruption. **f**, Increased F-actin (phalloidin in green) at the interface between basal and spinous cells in control and K10-Spastin epidermis. Arrow indicates the interface between basal and suprabasal cells. Scale bar, 20 μm. Quantification of interface F-actin intensity is shown below. In the K10-Spastin epidermis, only the interface F-actin between HA+ spinous and basal cells was measured. Data shown as the mean ± SEM, n=61 cells for control and n=86 cells for K10-Spastin from 3 embryos, p-value <0.0001, two-tailed unpaired t-test. **g**, Western blot of Myosin IIC and GAPDH in K10-Spastin and control epidermis at E16.5. **h**, Immunostaining of Myosin IIA in K10-Spastin and control epidermis at E16.5. Scale bar, 20 μm. **i**, Immunostaining of Myosin IIB in K10-Spastin and control epidermis at E16.5. Scale bar, 20 μm.

**Extended Data Fig. 2:**
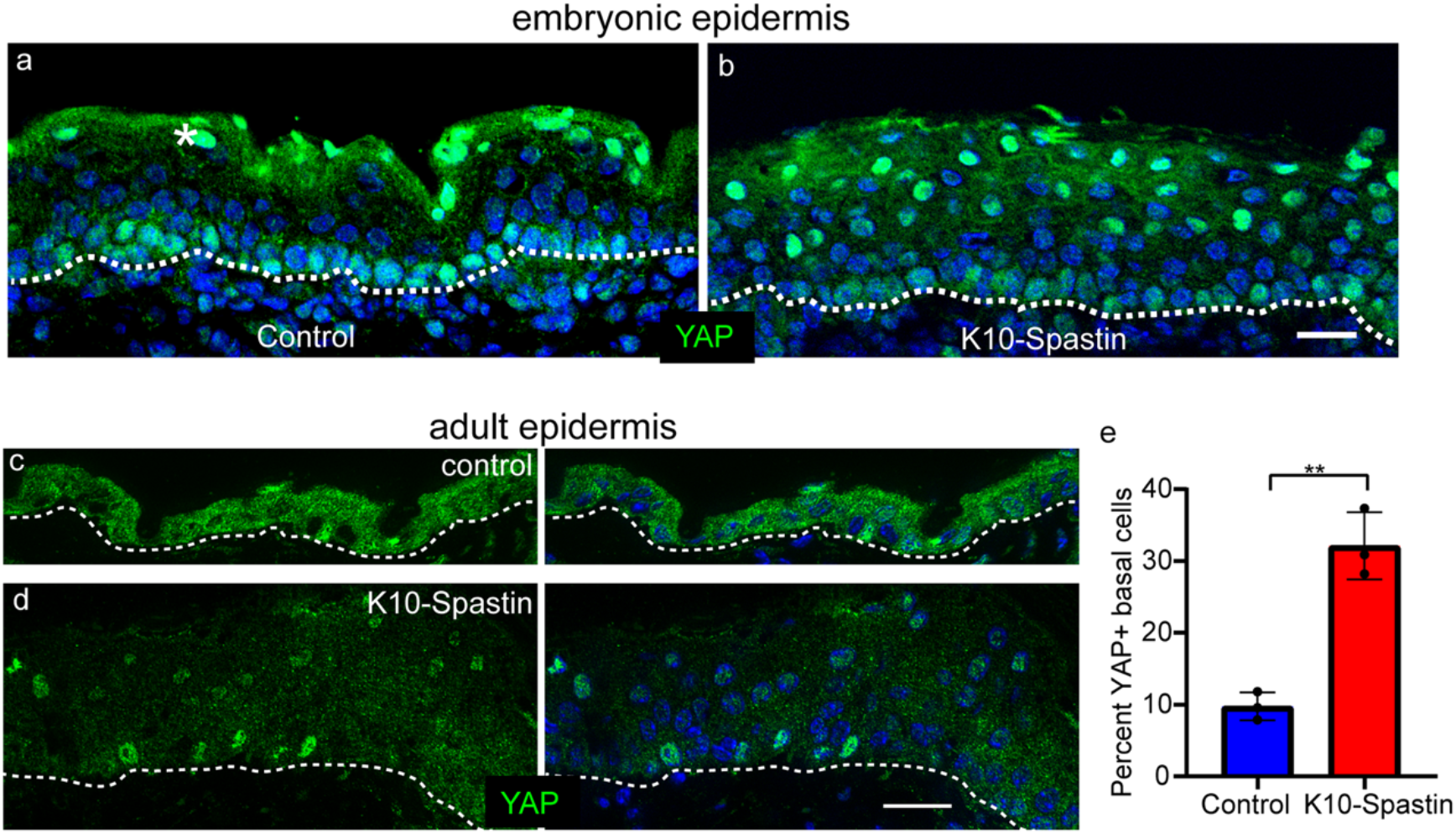
Microtubule depolymerization increases nuclear YAP in the epidermis, related to Fig. 2. **a,b**, Immunofluorescence staining of YAP (green) in control and K10-Spastin epidermis. This antibody showed non-specific labeling in the periderm. Notice the increased nuclear YAP signaling in the suprabasal cells in K10-Spastin. Scale bar, 20 μm. **c,d**, Immunostaining of YAP (green) in adult skin epidermis of control and K10-Spastin (on doxycycline for two weeks). **e**, Quantification of the percent of YAP positive basal cells in control and K10-Spastin. Data is shown as the mean +/− SD. N=3 mice for control (27 fields) and K10-Spastin (28 fields) were measured, p-value<0.01, two-tailed unpaired t-test.

**Extended Data Fig. 3:**
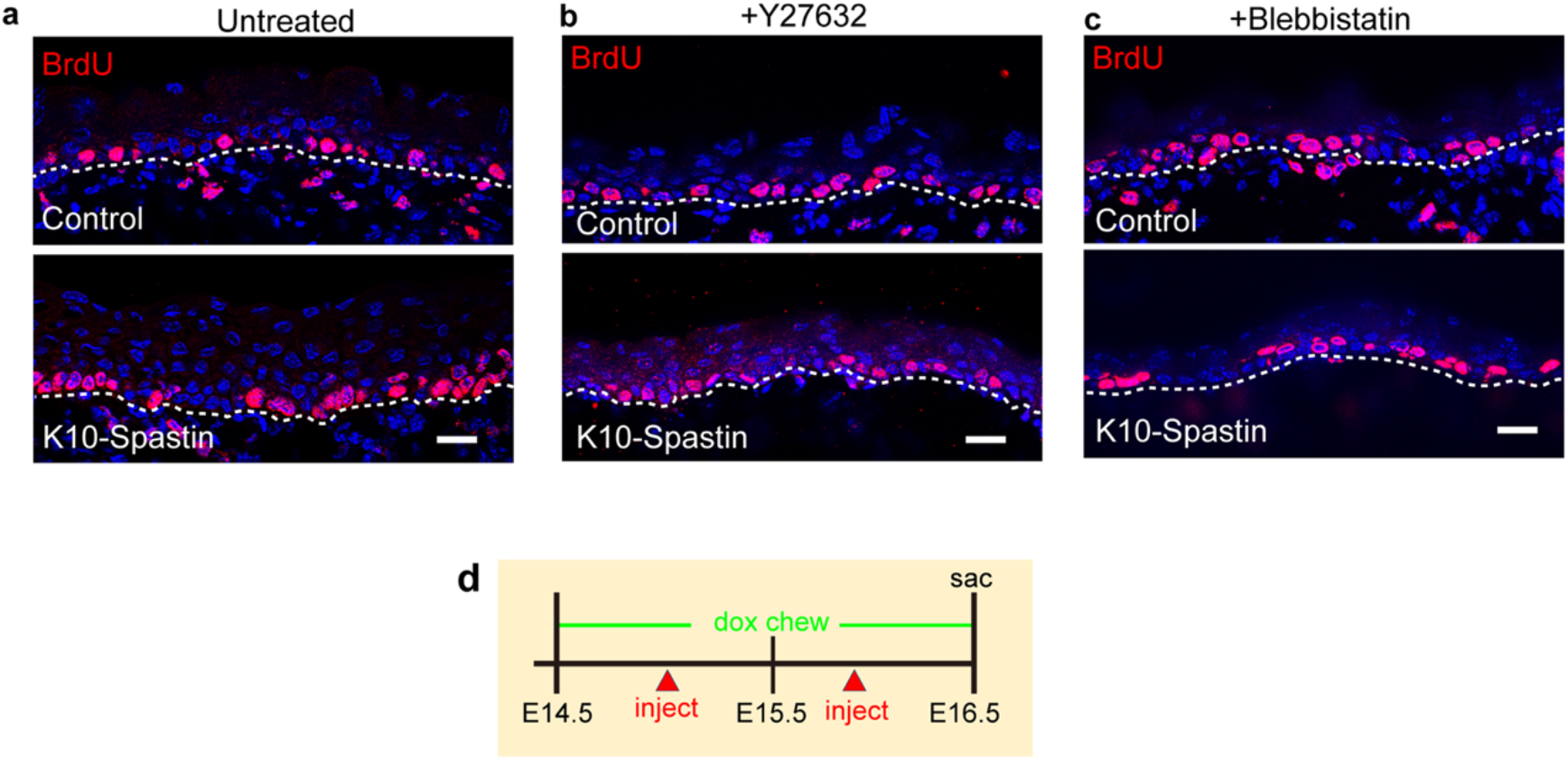
Inhibiting contractility rescues the basal cell hyperproliferation caused by microtubule depolymerization, related to Fig. 3. **a**, **b**, **c**, Immunofluorescence staining of BrdU in the untreated, Y27632, and Blebbistatin injected control and K10-Spastin epidermis respectively. Scale bars, 20 μm. **d**, Diagram depicting the drug injection times in K10-Spastin. Pregnant dams were fed with doxycycline chow from E14.5, and injected with these inhibitors twice (one early in the morning and one at night) the next day, and injected with BrdU 1 hour before sacrifice at E16.5.

**Extended Data Fig. 4:**
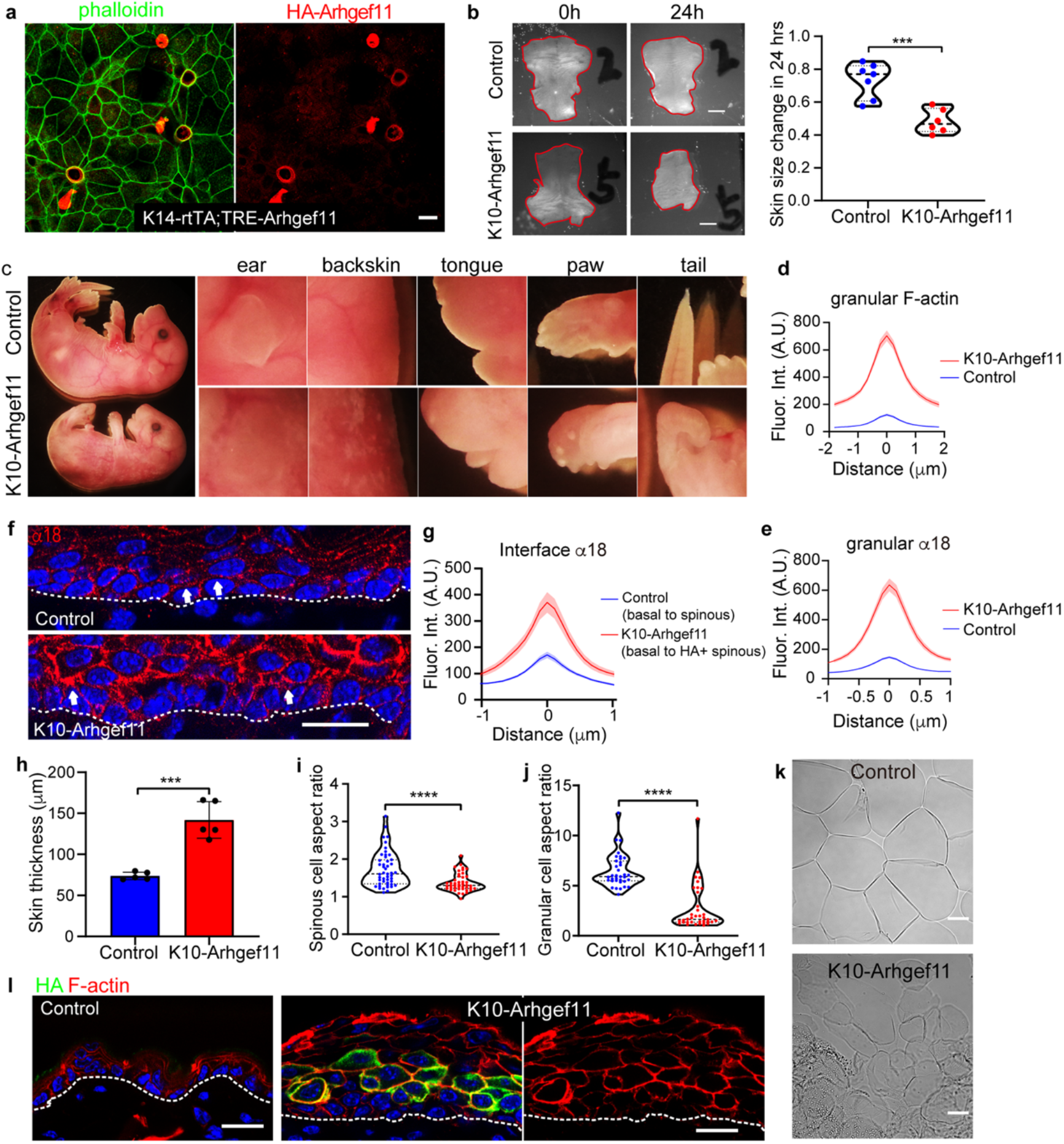
Increasing contractility by expressing Arhgef11^CA^ in differentiated cells induces skin thickening and cell rounding, related to Fig. 4. **a**, Immunostaining of F-actin (phalloidin, green) and HA-Arhgef11 (red) in keratinocytes transduced with K14-rtTA and pTRE2-Arhgef11 plasmids. These cells were treated with doxycycline and also with calcium to induce junction formation. Scale bar, 20 μm. **b**, **c**, Arhgef11^CA^ expression in differentiated cells of the epidermis leads to thick flaky skin, lack of external ears, mis-shapen paws, exposed tongue, and curly tails. **d**, Quantification of cortical F-actin fluorescence intensity in control and K10-Arhgef11granular cells. Data shown as the mean ± sem, n=26 cells for control and n=32 cells for K10-Arhgef11 from 3 embryos, p-value <0.0001, two-tailed unpaired t-test. **e**, Quantification of cortical α-18 fluorescence intensity in control and K10-Arhgef11granular cells. Data shown as the mean ± SEM, n=32 cells for control and n=34 cells for K10-Arhgef11 from 3 embryos, p-value <0.0001, two-tailed unpaired t-test. **f**, Increased α-18 at the interface between basal and spinous cells in control and K10-Arhgef11 at E17.5. Arrow indicates interface α-18. Scale bar, 20 μm. **g**, Quantification of interface α-18 intensity. In K10-Arhgef11, only F-actin at the interface between HA+ spinous and basal cells was measured. Data shown as the mean ± SEM, n=26 cells for control and n=23 cells for K10-Arhgef11 from 3 embryos, p-value <0.0001, two-tailed unpaired t-test. **h**, Quantification of skin thickness in K10-Arhgef11 and control at E17.5. Data is shown as the mean ± SD, n=5 embryos for control (489 regions) and K10-Arhgef11 (304 regions), p-value<0.001, two-tailed unpaired t-test. **i-j**, Quantification of cell aspect ratio in spinous and granular cells in K10-Arhgef11 and control respectively. For spinous cell quantification, n=3 embryos for control (47 cells) and K10-Arhgef11 (44 cells). For granular cells quantification, n=3 embryos for control (35 cells) and K10-Arhgef11 (36 cells). Data is shown as the mean ± SD, p-value<0.0001, two-tailed unpaired t-test. **k**, Isolated corneocytes from control and K10-Arhgef11 neonates at P0. Notice the abnormal morphology in K10-Arhgef11. Scale bar, 20 μm. **l**, Increased skin thickness and cortical F-actin (phalloidin in red) in Arhgef11^CA^ expressing cells (HA in green) compared to control at P37. Expression was induced at P29. Scale bar, 20 μm.

**Extended Data Fig. 5:**
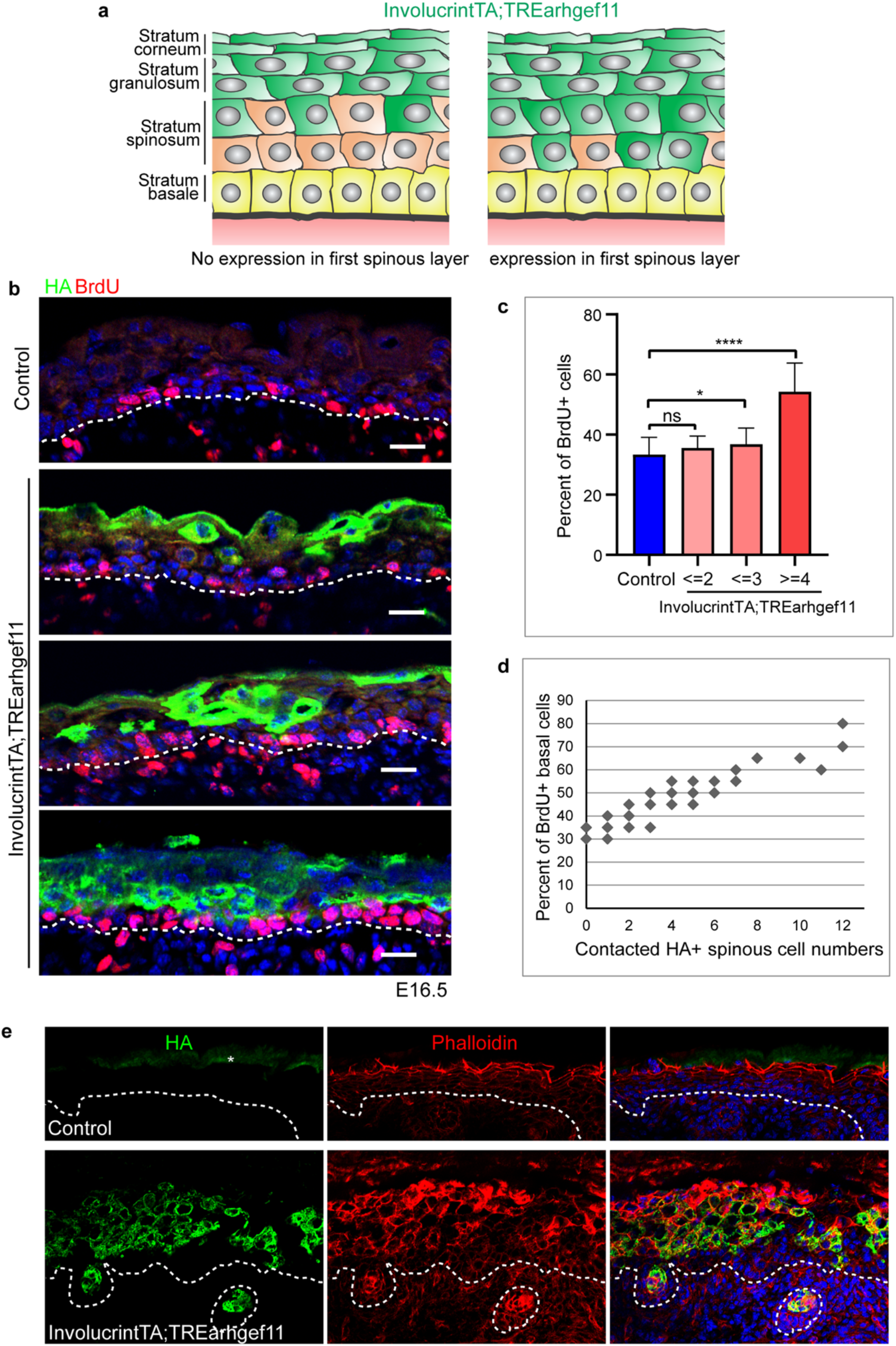
Increased contractility of cells not directly in contact with stem cells does not cause increased proliferation. **a**, Diagram depicting mosaic expression throughout the epidermis in Involucrint-TA;TRE-Arhgef11. **b**, BrdU (red) and HA (green) immunofluorescence staining in Involucrin-tTA;TRE-Arhgef11 and control at E16.5, doxycycline chow was removed at E12.5 to induce expression. Notice that BrdU incorporation in stem cells correlated with the number of contacted spinous cells expressing Arhgef11^CA^. Scale bars, 20 μm. **c**, Quantification of BrdU+ basal cells in control and Involucrin-tTA;TRE-Arhgef11. We quantitated random regions of epidermis that spanned 20 basal cells. In these regions, the number of BrdU+ and basal cell-contacted HA+ spinous cells were quantified. Notice that the percent of BrdU+ cells is significantly increased when there were over 3 spinous cells expressing Arhgef11^CA^. n=34 regions for control from 3 embryos and n=52 regions from 4 embryos. For analysis of control to (contacted HA+ spinous <=2), p-value= 0.1001, not significant; control to (contacted HA+ spinous<=3), p-value= 0.0172 (<0.05, significant); control to (contacted HA+ spinous>=4), p-value <0.0001. Data is shown as the mean ± SD, two-tailed unpaired t-test. **d**, Quantitation of BrdU+ basal cells with increasing numbers of spinous cells expressing Arhgef11^CA^. **e**, F-actin (phalloidin in red) and HA (green) immunofluorescence staining in control and Involucrin-tTA;TRE-Arhgef11 at E18.5 (doxycycline chow was removed at E12.5 to induce expression). Notice more robust expression of Arhgef11^CA^ in spinous cells and hair follicles at this later developmental stage, Scale bar, 40 μm.

**Extended Data Fig. 6:**
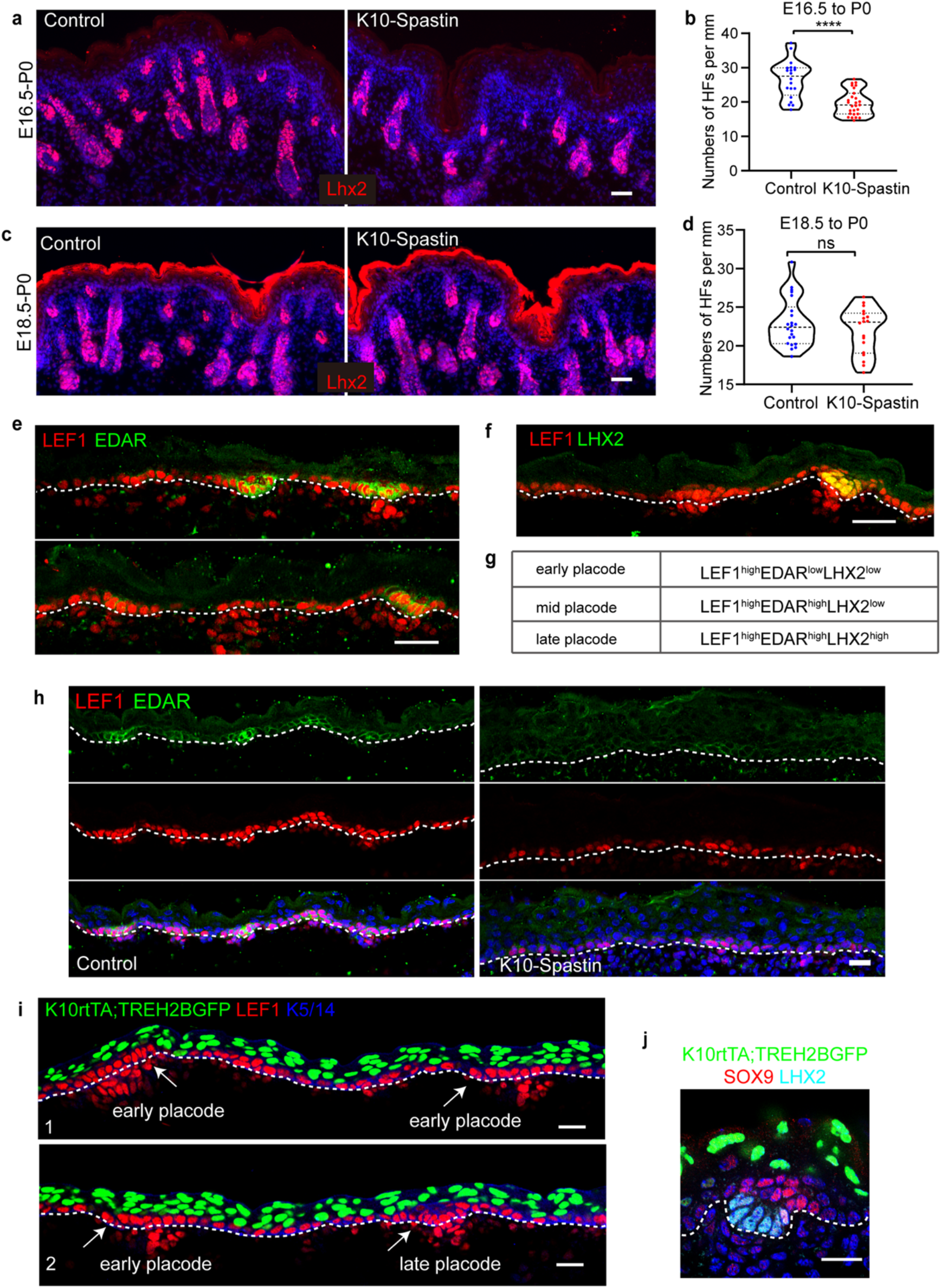
Increasing contractility in differentiated cells causes non-cell autonomous defects in hair follicle specification, related to Fig. 5 and Fig. 6. **a**-**b**, Staining of Lhx2 (red) of P0 epidermis in control and K10-Spastin (doxycycline treatment began at E16.5). Notice the normal development of the HFs that are already specified. Scale bar, 50 μm. Data shown as the mean ± SD, n=20 fields for control and n=27 fields for K10-Spastin from 2 embryos were measured, p-value<0.0001, two-tailed unpaired t-test. **c**-**d**, Staining of Lhx2 (red) in control and K10-Spastin skin P0 (doxycycline treatment began at E18.5). Scale bar, 50 μm. Data shown as the mean ± SD, n=22 fields for control and n=18 fields for K10-Spastin from 2 embryos were measured, p-value= 0.3023, not significant, two-tailed unpaired t-test. **e**, Staining of Lef1 (red) and EDAR (green) in control skin at E16.5. Notice the LEF1^high^ but EDAR^low^ early placode. Lef1 labels both the placode and dermal condensates below them. Scale bar, 30 μm. **f**, Staining of Lef1(red) and Lhx2 (green) in control skin at E16.5. Notice the Lef1^high^ but Lhx2^low^ early placode; Lhx2 labels only late stage placodes. Scale bar, 30 μm. **g**, Criteria for labeling placodes as early, mid and late. **h**, Co-staining of EDAR and Lef1 in control and K10-Spastin skin at E16.5, related to Fig. 6 f-g. Scale bar, 20 μm. **i**, Staining of Lef1 (red) and Krt5/14 (blue, basal layer marker) in K10-rtTA;TRE-H2B-GFP skin at E16.5. Note that there are no H2B-GFP labeled cells in either the early or late placode. Scale bar, 20 μm. **j**, Staining of Sox9 (red) and Lhx2 (cyan) in K10-rtTA;TRE-H2B-GFP skin at E16.5 showing no H2B-GFP induction in Sox9+ suprabasal cells in the late placode. Scale bar, 20 μm.

**Extended Data Fig. 7:**
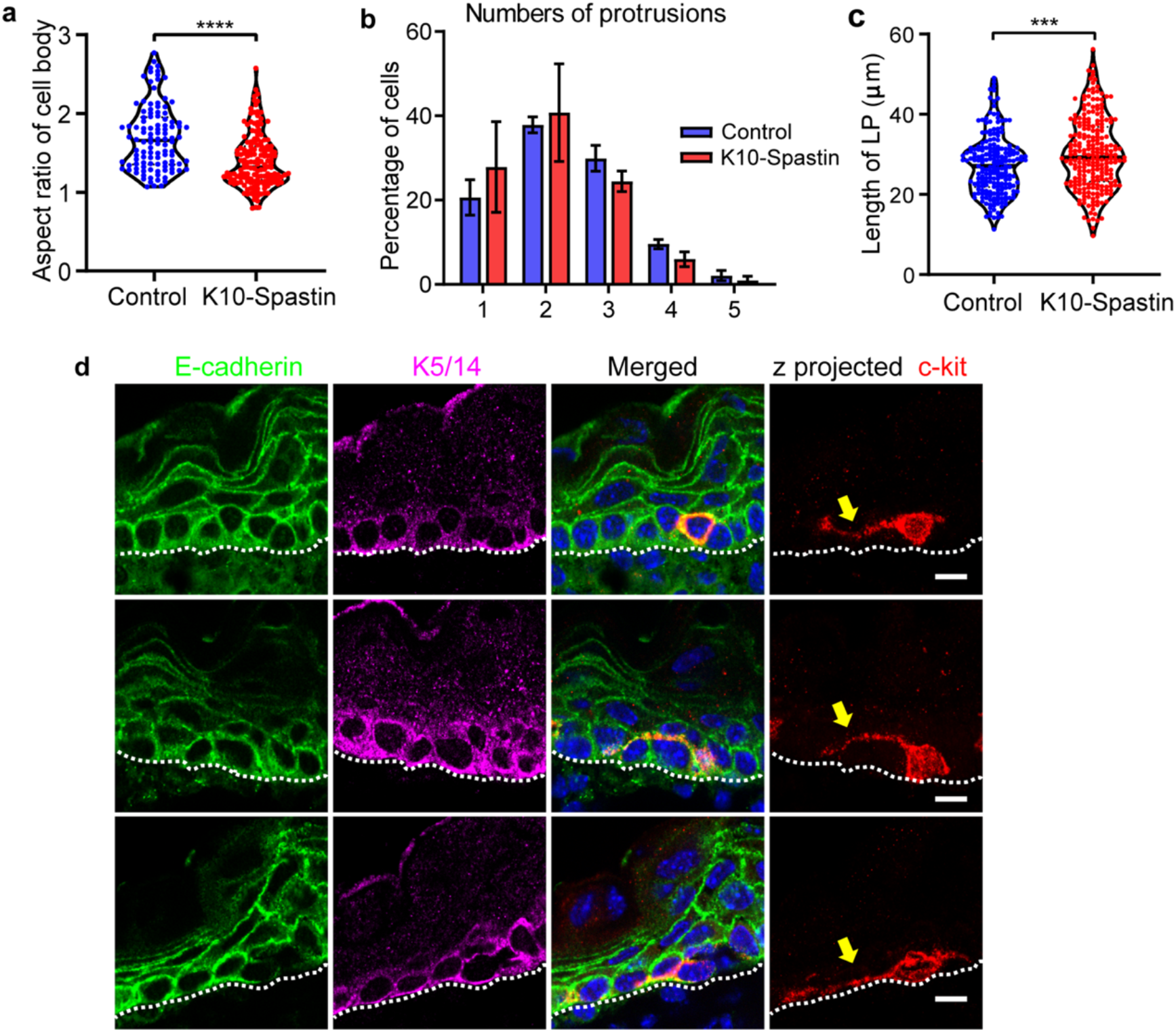
Increasing contractility in differentiated cells decreases migration of melanoblasts, related to Fig. 7. **a**, Quantification of cell body aspect ratio of melanoblasts in control and K10-Spastin epidermis. Data shown as the mean ± SD, n=110 melanoblasts for control and n=156 melanoblasts for K10-Spastin from 3 embryos, p-value<0.0001, two-tailed unpaired t-test. **b**, Quantification of protrusion numbers of melanoblasts in control and K10-Spastin. Data shown as the mean ± SD, n=3 embryos for control (320 melanoblasts) and for K10-Spastin (311 melanoblasts), p-value>0.9999, not significant, two-tailed paired t-test. **c**, Length of the leading protrusion in melanoblasts in control and K10-Spastin epidermis. Data shown as the mean ± SD, n=205 melanoblasts for control and n=237 melanoblasts for K10-Spastin from 3 embryos, p-value<0.001, two-tailed unpaired t-test. **d**, Co-staining of E-cadherin (green), the melanoblast marker C-kit (red) and the basal cell marker (Krt5/14) in control epidermis at E16.5. Arrows indicate the protrusions. Notice melanoblasts form protrusions that traverse through lateral junctions, above the basement membrane, and at the interface between suprabasal cells and basal cells. Scale bar, 10 μm.

**Extended Data Fig. 8:**
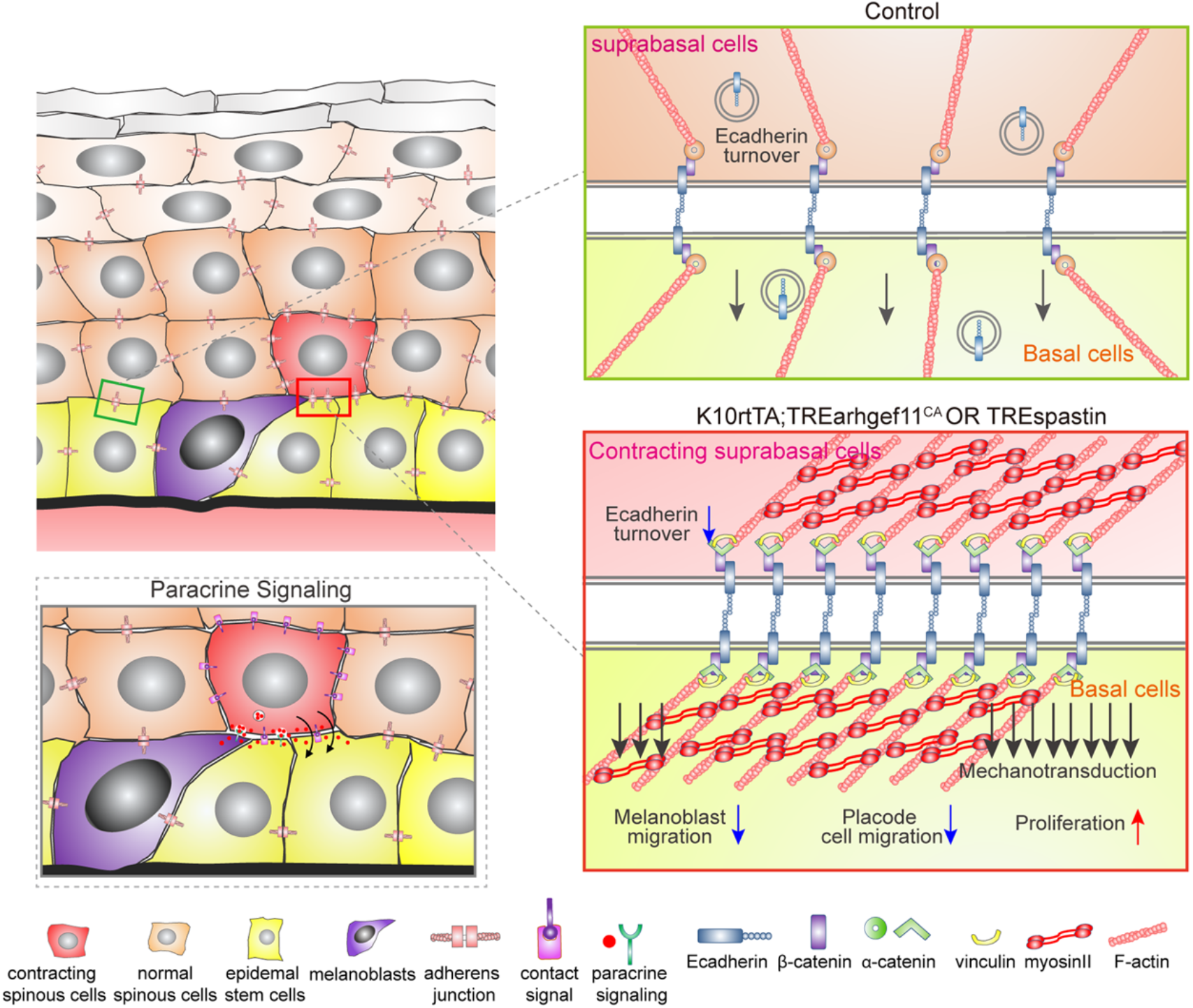
Model of regulation of basal stem cell proliferation and migration by their differentiated progeny. Our data demonstrates that differentiated suprabasal cells form part of the niche that controls the behavior of the underlying stem cells. Under normal conditions, basal stem cells are adhered to suprabasal spinous cells through adherens junction (and other structures) that are under low tension and rapid turnover. Increased contractility results in increased proliferation and decreased migration of underlying cells. There are two general models to explain this. First (lower left), contractility of daughter cells could alter their cell surface composition or their secretome, allowing them to signal to basal cells in a paracrine manner. A second possibility, which is not mutually exclusive, is that mechanical coupling through adherens junctions drives these phenotypes. In this case, either the increased stability of adherens junctions, or the increase tension across them would be the signal that inhibits migration and/or promotes proliferation.

## References

1. Fuchs, E., Tumbar, T. & Guasch, G. Socializing with the neighbors: stem cells and their niche. Cell 116, 769–778 (2004).

2. Cetera, M., Leybova, L., Joyce, B. & Devenport, D. Counter-rotational cell flows drive morphological and cell fate asymmetries in mammalian hair follicles. Nat. Cell Biol. 20, 541–552 (2018).

3. Ito, M. et al. Wnt-dependent de novo hair follicle regeneration in adult mouse skin after wounding. Nature 447, 316–320 (2007).

4. Ito, M. et al. Stem cells in the hair follicle bulge contribute to wound repair but not to homeostasis of the epidermis. Nat. Med. 11, 1351–1354 (2005).

5. Hsu, Y.C., Li, L. & Fuchs, E. Emerging interactions between skin stem cells and their niches. Nat. Med. 20, 847–856 (2014).

6. Watt, F.M. & Fujiwara, H. Cell-extracellular matrix interactions in normal and diseased skin. Cold Spring Harb. Perspect. Biol. 3, a005124 (2011).

7. Goldstein, J. & Horsley, V. Home sweet home: skin stem cell niches. Cell. Mol. Life Sci. 69, 2573–2582 (2012).

8. Sharp, L.L., Jameson, J.M., Cauvi, G. & Havran, W.L. Dendritic epidermal T cells regulate skin homeostasis through local production of insulin-like growth factor 1. Nat. Immunol. 6, 73–79 (2005).

9. Gur-Cohen, S. et al. Stem cell-driven lymphatic remodeling coordinates tissue regeneration. Science 366, 1218–1225 (2019).

10. Festa, E. et al. Adipocyte lineage cells contribute to the skin stem cell niche to drive hair cycling. Cell 146, 761–771 (2011).

11. Chen, D., Jarrell, A., Guo, C., Lang, R. & Atit, R. Dermal beta-catenin activity in response to epidermal Wnt ligands is required for fibroblast proliferation and hair follicle initiation. Development 139, 1522–1533 (2012).

12. Ali, N. et al. Regulatory T cells in skin facilitate epithelial stem cell differentiation. Cell 169, 1119–1129 (2017).

13. Watt, F.M. & Huck, W.T. Role of the extracellular matrix in regulating stem cell fate. Nat. Rev. Mol. Cell Biol. 14, 467–473 (2013).

14. Fuchs, E. & Raghavan, S. Getting under the skin of epidermal morphogenesis. Nat. Rev. Genet. 3, 199–209 (2002).

15. Jones, P.H., Harper, S. & Watt, F.M. Stem cell patterning and fate in human epidermis. Cell 80, 83–93 (1995).

16. Adams, J.C. & Watt, F.M. Changes in keratinocyte adhesion during terminal differentiation: reduction in fibronectin binding precedes alpha 5 beta 1 integrin loss from the cell surface. Cell 63, 425–435 (1990).

17. Watt, F.M. Role of integrins in regulating epidermal adhesion, growth and differentiation. EMBO J. 21, 3919–3926 (2002).

18. Grose, R. et al. A crucial role of beta 1 integrins for keratinocyte migration in vitro and during cutaneous wound repair. Development 129, 2303–2315 (2002).

19. Raghavan, S., Bauer, C., Mundschau, G., Li, Q. & Fuchs, E. Conditional ablation of beta1 integrin in skin. Severe defects in epidermal proliferation, basement membrane formation, and hair follicle invagination. J. Cell Biol. 150, 1149–1160 (2000).

20. Samuel, M.S. et al. Actomyosin-mediated cellular tension drives increased tissue stiffness and beta-catenin activation to induce epidermal hyperplasia and tumor growth. Cancer Cell 19, 776–791 (2011).

21. Discher, D.E., Janmey, P. & Wang, Y.L. Tissue cells feel and respond to the stiffness of their substrate. Science 310, 1139–1143 (2005).

22. Wang, Y., Wang, G., Luo, X., Qiu, J. & Tang, C. Substrate stiffness regulates the proliferation, migration, and differentiation of epidermal cells. Burns 38, 414–420 (2012).

23. Handorf, A.M., Zhou, Y., Halanski, M.A. & Li, W.J. Tissue stiffness dictates development, homeostasis, and disease progression. Organogenesis 11, 1–15 (2015).

24. Klein, E.A. et al. Cell-cycle control by physiological matrix elasticity and in vivo tissue stiffening. Curr. Biol. 19, 1511–1518 (2009).

25. Kechagia, J.Z., Ivaska, J. & Roca-Cusachs, P. Integrins as biomechanical sensors of the microenvironment. Nat. Rev. Mol. Cell Biol. 20, 457–473 (2019).

26. Tata, P.R. & Rajagopal, J. Regulatory Circuits and Bi-directional Signaling between Stem Cells and Their Progeny. Cell stem cell 19, 686–689 (2016).

27. Hsu, Y.C. & Fuchs, E. A family business: stem cell progeny join the niche to regulate homeostasis. Nat. Rev. Mol. Cell Biol. 13, 103–114 (2012).

28. Guo, Z. & Ohlstein, B. Stem cell regulation. Bidirectional Notch signaling regulates Drosophila intestinal stem cell multipotency. Science 350(2015).

29. Jiang, H. et al. Cytokine/Jak/Stat signaling mediates regeneration and homeostasis in the Drosophila midgut. Cell 137, 1343–1355 (2009).

30. Amcheslavsky, A., Jiang, J. & Ip, Y.T. Tissue damage-induced intestinal stem cell division in Drosophila. Cell stem cell 4, 49–61 (2009).

31. Liang, J., Balachandra, S., Ngo, S. & O’Brien, L.E. Feedback regulation of steady-state epithelial turnover and organ size. Nature 548, 588–591 (2017).

32. Mondal, B.C. et al. Interaction between differentiating cell- and niche-derived signals in hematopoietic progenitor maintenance. Cell 147, 1589–1600 (2011).

33. Tadokoro, T., Gao, X., Hong, C.C., Hotten, D. & Hogan, B.L. BMP signaling and cellular dynamics during regeneration of airway epithelium from basal progenitors. Development 143, 764–773 (2016).

34. Muroyama, A. & Lechler, T. A transgenic toolkit for visualizing and perturbing microtubules reveals unexpected functions in the epidermis. eLife 6, e29834 (2017).

35. Zaidel-Bar, R., Zhenhuan, G. & Luxenburg, C. The contractome--a systems view of actomyosin contractility in non-muscle cells. J. Cell Sci. 128, 2209–2217 (2015).

36. Vicente-Manzanares, M., Ma, X., Adelstein, R.S. & Horwitz, A.R. Non-muscle myosin II takes centre stage in cell adhesion and migration. Nat. Rev. Mol. Cell Biol. 10, 778–790 (2009).

37. le Duc, Q. et al. Vinculin potentiates E-cadherin mechanosensing and is recruited to actin-anchored sites within adherens junctions in a myosin II-dependent manner. J. Cell Biol. 189, 1107–1115 (2010).

38. Yonemura, S., Wada, Y., Watanabe, T., Nagafuchi, A. & Shibata, M. alpha-Catenin as a tension transducer that induces adherens junction development. Nat. Cell Biol. 12, 533–542 (2010).

39. Zhang, H., Pasolli, H.A. & Fuchs, E. Yes-associated protein (YAP) transcriptional coactivator functions in balancing growth and differentiation in skin. Proc. Natl. Acad. Sci. U.S.A. 108, 2270–2275 (2011).

40. Rao, M.V. & Zaidel-Bar, R. Formin-mediated actin polymerization at cell-cell junctions stabilizes E-cadherin and maintains monolayer integrity during wound repair. Mol. Biol. Cell 27, 2844–2856 (2016).

41. Verma, S. et al. A WAVE2-Arp2/3 actin nucleator apparatus supports junctional tension at the epithelial zonula adherens. Mol. Biol. Cell 23, 4601–4610 (2012).

42. Snippert, H.J. et al. Intestinal crypt homeostasis results from neutral competition between symmetrically dividing Lgr5 stem cells. Cell 143, 134–144 (2010).

43. Kovacs, M., Toth, J., Hetenyi, C., Malnasi-Csizmadia, A. & Sellers, J.R. Mechanism of blebbistatin inhibition of myosin II. J. Biol. Chem. 279, 35557–35563 (2004).

44. Pellegrin, S. & Mellor, H. Actin stress fibres. J. Cell Sci. 120, 3491–3499 (2007).

45. Valon, L., Marin-Llaurado, A., Wyatt, T., Charras, G. & Trepat, X. Optogenetic control of cellular forces and mechanotransduction. Nat. Commun. 8, 14396 (2017).

46. Jaubert, J., Patel, S., Cheng, J. & Segre, J.A. Tetracycline-regulated transactivators driven by the involucrin promoter to achieve epidermal conditional gene expression. J. Invest. Dermatol. 123, 313–318 (2004).

47. Tsai, S.Y. et al. Wnt/beta-catenin signaling in dermal condensates is required for hair follicle formation. Dev. Biol. 385, 179–188 (2014).

48. Andl, T., Reddy, S.T., Gaddapara, T. & Millar, S.E. WNT signals are required for the initiation of hair follicle development. Dev. Cell 2, 643–653 (2002).

49. Ouspenskaia, T., Matos, I., Mertz, A.F., Fiore, V.F. & Fuchs, E. WNT-SHH Antagonism Specifies and Expands Stem Cells prior to Niche Formation. Cell 164, 156–169 (2016).

50. Zhang, Y. et al. Reciprocal requirements for EDA/EDAR/NF-kappaB and Wnt/beta-catenin signaling pathways in hair follicle induction. Dev. Cell 17, 49–61 (2009).

51. Fu, J. & Hsu, W. Epidermal Wnt controls hair follicle induction by orchestrating dynamic signaling crosstalk between the epidermis and dermis. J. Invest. Dermatol. 133, 890–898 (2013).

52. Tomann, P., Paus, R., Millar, S.E., Scheidereit, C. & Schmidt-Ullrich, R. Lhx2 is a direct NF-kappaB target gene that promotes primary hair follicle placode down-growth. Development 143, 1512–1522 (2016).

53. Ahtiainen, L. et al. Directional cell migration, but not proliferation, drives hair placode morphogenesis. Dev. Cell 28, 588–602 (2014).

54. Mort, R.L., Jackson, I.J. & Patton, E.E. The melanocyte lineage in development and disease. Development 142, 620–632 (2015).

55. Dogterom, M. & Koenderink, G.H. Actin-microtubule crosstalk in cell biology. Nat. Rev. Mol. Cell Biol. 20, 38–54 (2019).

56. Rodriguez, O.C. et al. Conserved microtubule-actin interactions in cell movement and morphogenesis. Nat. Cell Biol. 5, 599–609 (2003).

57. Krendel, M., Zenke, F.T. & Bokoch, G.M. Nucleotide exchange factor GEF-H1 mediates cross-talk between microtubules and the actin cytoskeleton. Nat. Cell Biol. 4, 294–301 (2002).

58. Nagae, S., Meng, W. & Takeichi, M. Non-centrosomal microtubules regulate F-actin organization through the suppression of GEF-H1 activity. Genes Cells 18, 387–396 (2013).

59. Muroyama, A., Terwilliger, M., Dong, B., Suh, H. & Lechler, T. Genetically induced microtubule disruption in the mouse intestine impairs intracellular organization and transport. Mol. Biol. Cell 29, 1533–1541 (2018).

60. Gasic, I., Boswell, S.A. & Mitchison, T.J. Tubulin mRNA stability is sensitive to change in microtubule dynamics caused by multiple physiological and toxic cues. PLoS Biol. 17, e3000225 (2019).

61. Benham-Pyle, B.W., Pruitt, B.L. & Nelson, W.J. Cell adhesion. Mechanical strain induces E-cadherin-dependent Yap1 and beta-catenin activation to drive cell cycle entry. Science 348, 1024–1027 (2015).

62. Indra, I., Troyanovsky, R.B., Shapiro, L., Honig, B. & Troyanovsky, S.M. Sensing Actin Dynamics through Adherens Junctions. Cell Rep. 30, 2820–2833 (2020).

63. Chrostek, A. et al. Rac1 is crucial for hair follicle integrity but is not essential for maintenance of the epidermis. Mol. Cell. Biol. 26, 6957–6970 (2006).

64. Jackson, B. et al. RhoA is dispensable for skin development, but crucial for contraction and directed migration of keratinocytes. Mol. Biol. Cell 22, 593–605 (2011).

65. Sumigray, K.D., Foote, H.P. & Lechler, T. Noncentrosomal microtubules and type II myosins potentiate epidermal cell adhesion and barrier formation. J. Cell Biol. 199, 513–525 (2012).

66. Subramanian, A. et al. Gene set enrichment analysis: a knowledge-based approach for interpreting genome-wide expression profiles. Proc. Natl. Acad. Sci. U.S.A. 102, 15545–15550 (2005).

67. Wang, J., Vasaikar, S., Shi, Z., Greer, M. & Zhang, B. WebGestalt 2017: a more comprehensive, powerful, flexible and interactive gene set enrichment analysis toolkit. Nucleic Acids Res. 45, W130–W137 (2017).

68. Tinevez, J.Y. et al. TrackMate: An open and extensible platform for single-particle tracking. Methods 115, 80–90 (2017).

